# Representation of emotional expressions across the face and voice brain networks

**DOI:** 10.64898/2025.12.22.695403

**Authors:** Stefania Mattioni, Federica Falagiarda, Rémi Gau, Mohamed Rezk, Ceren Battal, Alice Van Audenhaege, Olivier Collignon

**Affiliations:** Department of Experimental Psychology & Center for Cognitive Neuroscience (CCN), University of Gent, Belgium; Institutes for research in Neuroscience (IoNS), University of Louvain (UCLouvain), Belgium; Faculty of Psychology and Educational Sciences, University of Coimbra, Portugal

## Abstract

Recognizing emotion expressions from facial and vocal signals is crucial for optimal social interactions. We comprehensively characterized regions within the face and voice processing networks to probe their contribution to unimodal and crossmodal emotion representation. Using multivariate pattern classification analyses, we found that emotional expressions could be reliably decoded from the dominant sensory modality in all individually defined face- and voice-selective regions. Emotion expressions from the non-dominant modality (vocal expressions in the face networks, and vice versa) could also be decoded in all areas except the occipital face areas. A shared neural code for facial and vocal emotions is implemented in the temporal voice areas (TVA), the posterior superior temporal sulcus (pSTS) and the precentral gyrus (PCG). The simultaneous presentation of congruent facial and vocal expressions elicited distinct activity patterns across most regions, highlighting that multisensory inputs reshape brain responses relative to unisensory stimulation across the entire face and voice brain network.

These findings suggest that face and voice-selective regions broadly encode emotion expressions within and even across the senses, relying on a multisensory gradient that converges in temporo-frontal regions where their distributed multisensory responses align to create a supramodal representation of specific emotion expressions.

## Introduction

A person’s emotional state can be readily inferred from facial or vocal expressions. Specific configurations of facial muscle movements (Ekman & Friesen, 1971) or acoustic utterances (Sauter & Eimer, 2010) broadcast a person’s emotional state, and the ability of humans and other animals to efficiently recognize these emotional expressions is arguably one of the most important perceptual skills for successful social interactions (Darwin, 1872). Face perception relies on neural computations distributed across a series of face-selective regions in the occipital and temporal cortex (Duchaine & Yovel, 2015; Kanwisher & Yovel, 2006; Pitcher, Walsh, et al., 2011). In humans, face-selective regions are thought to be organized into two processing streams (Finzi et al., 2021; Pitcher & Ungerleider, 2021; Weiner & Grill-Spector, 2013). The ventral face processing stream, which is primarily involved in face recognition, is thought to be mainly tuned to invariant face features (Andrews & Ewbank, 2004; Grill-Spector et al., 2004; Kanwisher et al., 1997; Winston et al., 2004) and chiefly includes the occipital face area (OFA) and the fusiform face area (FFA). Second, the lateral stream is thought to primarily include the posterior superior temporal cortex (pSTS) and is instead associated with the processing of dynamic (Duchaine & Yovel, 2015; Pitcher, Dilks, et al., 2011; Weiner & Grill-Spector, 2013), social (Andrews & Ewbank, 2004; Duchaine & Yovel, 2015; Pitcher & Ungerleider, 2021), and multimodal perception (Pitcher & Ungerleider, 2021; Zhu & Beauchamp, 2017), as it specializes in dynamically changing aspects of faces (Pitcher et al., 2014; Puce et al., 1998). This view predicts that the encoding of emotional expressions is restricted to the pSTS (Haxby et al., 2000; Pitcher & Ungerleider, 2021), an idea that is nevertheless challenged by studies reporting that emotional expressions also modulate the activity in the ventral face processing stream (Ganel et al., 2005; Harry et al., 2013; Kawasaki et al., 2012; Monroe et al., 2013; Vuilleumier & Pourtois, 2007; Wegrzyn et al., 2015)).

Voice perception also relies on specific voice-selective regions located in the temporal cortex, often referred to as the temporal voice areas (TVAs; Belin et al., 2000). Early accounts proposed that vocal emotion selectively engages regions beyond the TVAs, overlapping with face-selective areas in the pSTS (e.g., Ethofer et al., 2006; Grandjean et al., 2005; Kotz et al., 2013; Wildgruber et al., 2005). Similar to face processing, this view remains debated, as other studies have suggested that emotional information can be represented throughout the entire set of TVAs regions (Ethofer et al., 2009).

The way we express and perceive emotions is inherently multisensory. Emotion expressions are typically conveyed through multiple spatiotemporally congruent, modality-specific cues, together with a set of invariant, amodal features (Lewkowicz & Ghazanfar, 2009). Similar perceptual mechanisms are engaged when recognizing emotions from both faces and voices (Kuhn et al., 2017), and the ability to integrate multisensory emotional information from talking faces is crucial for extracting coherent meaning from these ubiquitous communicative signals (Ghazanfar & Logothetis, 2003; Lewkowicz & Ghazanfar, 2009). When facial and vocal expressions are presented together, we discriminate emotions better and faster (Collignon et al., 2010; Falagiarda & Collignon, 2019, p. 202; Kreifelts et al., 2007), and this integration appears to be automatic and independent of available attentional resources (Collignon et al., 2008; Vroomen et al., 2001). The integration of emotional expressions into a common representation likely relies on core evolutionary mechanism (Darwin, 1872), as non-human primates (Ghazanfar et al., 2005) and young infants (Walker-Andrews & Lennon, 1991) also seamlessly match the congruency of facial and vocal signals. However, which brain regions jointly represent and integrate facial and vocal emotional expressions remains poorly understood (Klasen et al., 2012; Young et al., 2020). Classical models of face and voice processing propose that “unisensory” nodes sensitive to facial and vocal emotional expressions transmit a categorical emotional readout signals to higher-order brain regions, such as the posterior superior temporal sulcus (pSTS) or the medial prefrontal cortex (MPFC), where a unified – abstract – representation of emotional expressions is formed (Davies-Thompson et al., 2019; Hagan et al., 2009; Kreifelts et al., 2007; Peelen et al., 2010; Watson et al., 2014). In contrast to this strictly hierarchical view—in which categories of emotional expressions extracted from faces and voices converge into a common neural representation—some studies point to the possibility of cross-sensory interactions between areas that are traditionally considered unimodal. Support for this idea comes notably from non-human primate studies showing that neurons in voice-selective regions alter their response to vocalizations in the presence of congruent or incongruent facial expressions (Ghazanfar et al., 2005; see also Dureux et al., 2024; Khandhadia et al., 2021 for similar observation in face patches of macaque). In humans, several studies have shown altered univariate responses in putatively “unimodal” regions (e.g. FFA) in response to crossmodal input (voice) during tasks requiring identity discrimination (Kriegstein et al., 2005; Lowe et al., 2021; Von Kriegstein & Giraud, 2006). Furthermore, diffusion imaging studies have suggested the existence of a direct structural connection between the FFA and the TVA, two key regions of the face and voice networks, respectively (Benetti et al., 2018; Blank et al., 2011). Building on this distributed perspective, a recent study found that the fusiform gyrus in the visual cortex encodes abstract emotion categories from both visual and auditory modalities (Lettieri et al., 2024).

A comprehensive investigation of how multisensory emotional expressions are represented across the face and voice processing networks is still lacking. Here, we address this gap by characterizing the brain activity elicited by visual, auditory, and bimodal emotional expressions from five distinct categories –neutral, disgust, fear, happiness, and sadness – in individually defined face- and voice-selective regions. Our goal was to address four main questions. First, we investigated where facial and vocal emotional expressions are represented within their respective networks. Second, we explored whether traces of crossmodal emotional information could be detected in the identified regions (i.e. vocal emotional information in the face network and facial emotional information in the voice network). Third, in regions jointly representing facial and vocal emotional expressions, we examined whether emotional expressions are represented, at least partially, independently of the sensory modality of the input. Fourth, we investigated which regions of the face and voice networks show distinct response patterns to multisensory compared to unisensory signals, assessing whether bimodal responses align with the dominant sensory modality or instead form a distinct bimodal representation. These questions were addressed using a series of multivariate decoding and representational similarity analyses applied to functional magnetic resonance imaging (fMRI) recordings.

## Methods

### Participants

Twenty-four (twelve male, age range: 20-44) neurotypically developed adults and naïve to the experimental hypothesis, took part in the study. All participants were native French speakers, right handed and had normal or corrected-to-normal vision (corrected in the scanner with MRI-compatible glasses when needed) and self-reported normal audition. They provided their written informed consent in the first experimental session, prior to the start of any data collection.

### Stimuli

The experiment employs 20 audio-visual clips as stimuli. Each video portrays one of four actors (two males, two females) expressing one of the five following expressions: disgusted, fearful, happy, sad or neutral. The clips last one second and aim at showing the emotional state through the facial expression as well as a concomitant non-linguistic vocalization. The stimuli were selected among a wider pool of actors and actresses (pre-selection pool of recordings, a subset of which can be found in Belin et al., 2008 (vocal component) and Simon et al., 2008 (visual component), which were rated, separately for the face and voice expressions, by an independent group (N=20). The chosen stimuli did not induce any specific discrimination bias in the five selected expressions, neither in vision nor in audition. In addition to the face and voice stimuli, twenty videos and twenty audio tracks of everyday objects and scenes were used in the face and voice localizer respectively (see section *Localizers*). These stimuli were taken from the internet and were copyright free.

The use of dynamic stimuli in the investigation of visual perception in general, and facial perception specifically, is crucial for recreating the vivid experience that we typically encounter in everyday life. It has been shown that, compared to static images, dynamic stimulation produces higher discrimination performance, differential neural activation and successful decoding through machine learning (Davies-Thompson et al., 2019; Grèzes et al., 2007; Liang et al., 2018; Trautmann-Lengsfeld et al., 2013). Dynamic visual stimuli are even more important when investigating audio-visual perception and integration, where the auditory component cannot be un-embedded from its time course, and where a static visual stimulation would always have a limited generalizability.

### Design and procedure

The experimental protocol was approved by the Ethical committee of the University Hospital of St Luc (Université Catholique de Louvain), Brussels, Belgium. The entirety of the experiment spanned two experimental sessions at St Luc Hospital, with a duration of about 2 hours each. The first session consisted in a brief phase of behavioral rating of the stimuli, followed by the MRI acquisition. The second session included only the MRI acquisition part. Each participant spent around 90 minutes in the scanner in each session, for a total of about 3 hours of MRI data acquisition for each participant.

In the two sessions inside the MRI scanner, each participant attended several acquisition runs. The runs included two block-design functional localizers and a total of 54 short event-related runs with three different modalities of stimuli presentation: visual only (facial expression), auditory only (non-linguistic vocalization) or bimodal stimulation (congruent facial and vocal expression). One high resolution structural image was also acquired in the first session for each subject (see Acquisition parameters).

In each session, the participant completed one of the two functional localizers as well as nine runs in each of the three modalities. Across the two sessions, 18 runs per subject per modality were acquired. Each participant additionally completed a short behavioral experiment, which took place on the first day of the fMRI experiment, before the commencing of any MRI data acquisition. During this behavioral session, participants were asked to rate each unimodal stimulus on several scales (see *Supplemental Material* for more details) divided into two tasks and blocked by modality (20 visual stimuli and 20 auditory stimuli rated independently, tasks’ order was counterbalanced across participants). These data are not analyzed or discussed in the current report, however it is relevant to note that participants were familiarized with the stimuli prior to the scanning, to ensure that the emotion expressed in the stimuli would be recognized as the emotion categories we expected.

#### Localizers

We used functional localizers for the definition of regions of interest in individual subjects. The aim of both face and voice localizers was to provide individually defined activation clusters of the core face and voice network regions. We were open to the inclusion of more areas that the localizers would consistently reveal across participants. The localizer designs (explained more in detail in the upcoming paragraphs), were initially piloted on six subjects that are not included in the final sample of participants.

#### Face localizer

The face localizer, which aims at the definition of the core face network in each participant, consists in one functional acquisition run with a block-design alternating ten blocks of faces to ten blocks of objects/scenes (see Figure 1A). Each block of either faces or objects consisted in the presentation of twenty 1s-long videos, with a 100ms inter-stimulus interval (ISI). The facial expressions videos were the same twenty clips used in the main experiment, while the twenty objects/scenes contained in the objects’ block were similar to the ones utilized in a well-established dynamic face localizer (Fox et al., 2009), but downloaded anew from the internet and trimmed to a 1s length when needed (see the Supplemental Material for the exhaustive list of stimuli). We included ten seconds of presentation of the background only, at the beginning, middle and end of the run, to be used to estimate the baseline. All videos subtended a frame of 720 x 480 pixels on the screen, which corresponded to 8.5 degrees horizontally and 5.7 degrees vertically, of visual angle at a distance of about 175cm.

**Figure 1.**
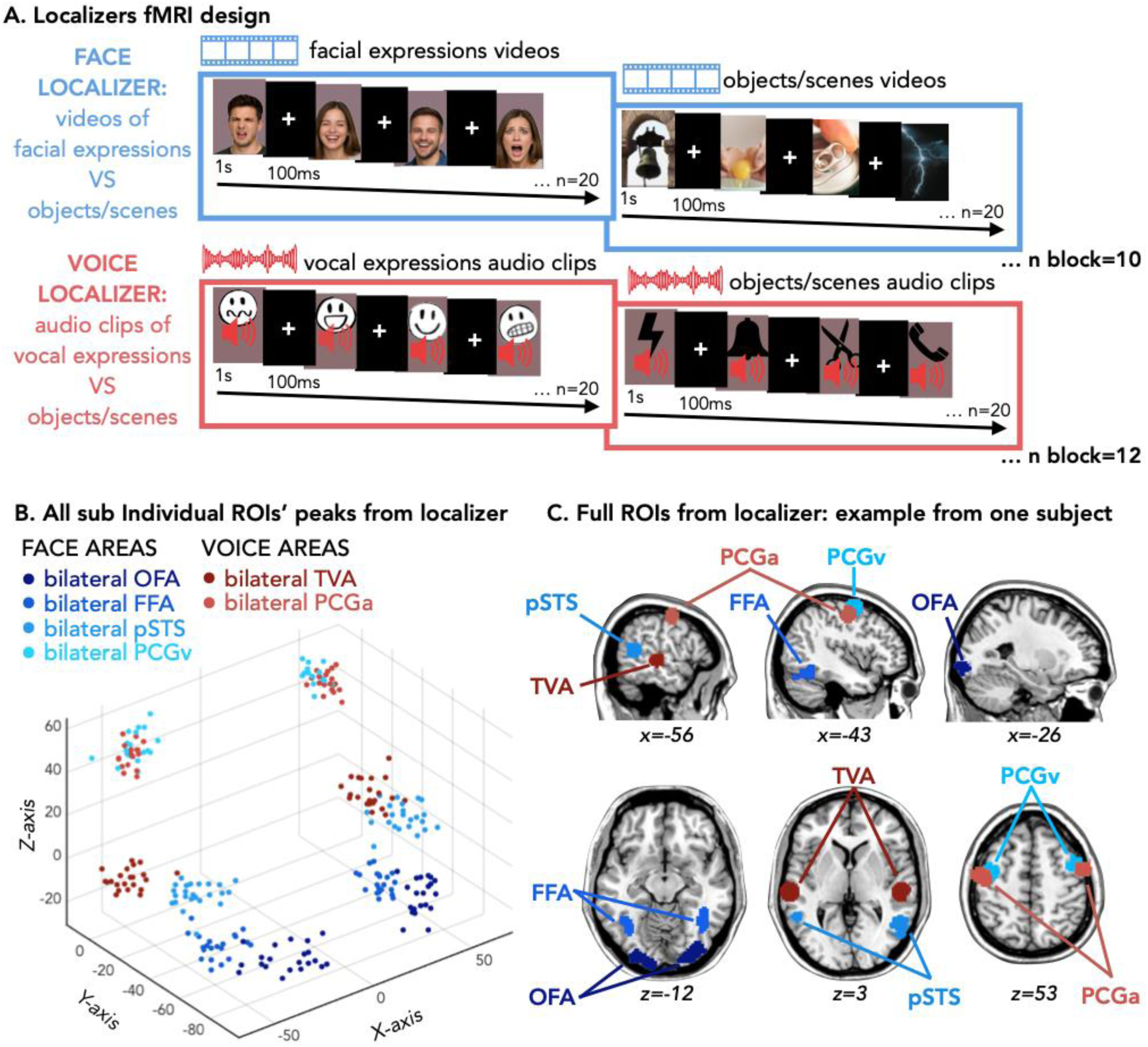
**A)** Experimental design implemented in the fMRI for the face (in blue) and voice (in red) localizers. The facial images shown in this figure are AI-generated and are used for illustrative purposes only; they are not screenshots extracted from the experimental video stimuli and do not depict real individuals. **B)** Results from the localizer, representing each individual peak for the contrasts [faces>objects] in dark to light blue and for [voices>objects] in dark to light red (see all individual peaks in Table S1 and Table S2). C) Example of the 12 full ROIs built around one subject’s individual peaks.

#### Voice localizer

Analogous to the face localizer, the voice localizer aims at the definition of the core voice network and consists in a functional acquisition with a block-design, alternating twelve blocks of voices to twelve blocks of auditory objects (see Figure 1A). Each block of either voices or objects consisted in the presentation of twenty 1s-long audio tracks, with 100ms ISI. All soundtracks were RMS-equalized and presented through in-ear MRI-compatible earphones (Sensimetric S14, SensiMetrics, USA). The vocal expressions were the same used in the main experiment, while the auditory objects’ sounds (see the Supplemental Material for the exhaustive list of stimuli) were downloaded from the internet and trimmed to a 1s length when needed (copyright free sounds from freesounds.org). We included ten seconds of silence at the beginning, middle and end of the run for baseline. Participants were asked to set a loud but comfortable volume for the auditory stimuli at the beginning of each scanning session.

#### Main experiment

In the main experiment, only stimuli coming from the audio-visual emotional clips were presented. The experiment, conducted over two sessions, consists of fifty-four runs divided into three modality conditions: eighteen runs for the visual, eighteen runs for the auditory, and eighteen runs for the bimodal tasks. All runs of the main experiment have an event-related design and every run contains one repetition of each stimulus, i.e. 20 stimuli. In the visual task, the rendered videos were cropped to 332 pixels wide and 424 pixels high, sustaining 3.9 degrees horizontally and 5 degrees vertically of visual angles in the participants’ visual field. For the auditory task, the audio track was extracted from the original clips as a wav file and presented through in-ear MRI-compatible earphones (Sensimetric S14, SensiMetrics, USA). During the auditory presentation, participants were asked to keep their eyes open and looked at a fixation cross that appeared on screen during the event. In the bimodal task, both the audio track and the video frames were presented and rendered as a synchronized, always congruent, audio-visual clip. In all tasks, each stimulus was presented twice in a row, for an event duration of 2 seconds (2 x 1-second file) and by an inter-stimulus interval (ISI) of 3 seconds (see Figure 2A for illustration of trial sequences of the main experimental runs).

**Figure 2.**
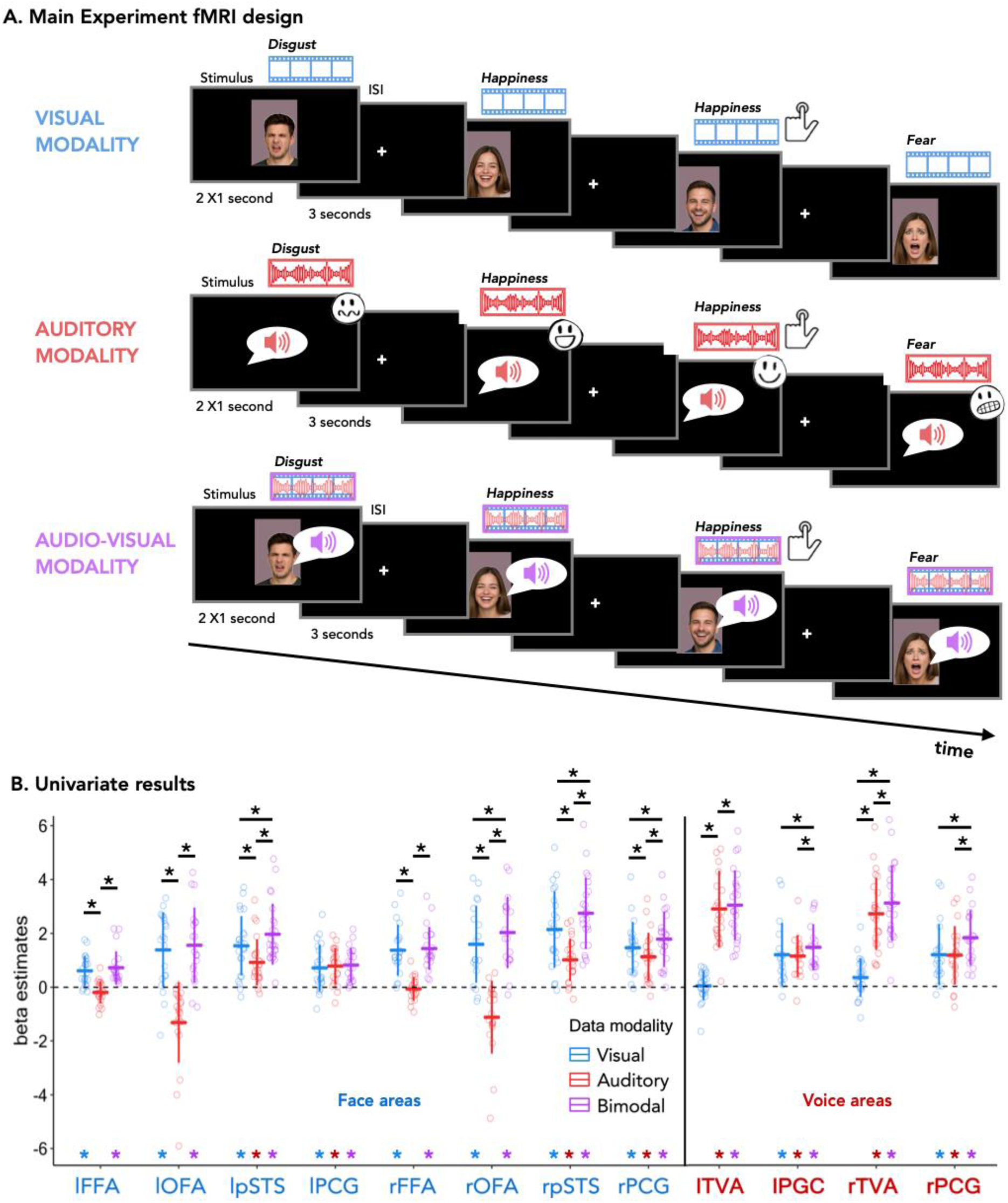
**A)** Experimental design implemented in the fMRI for the main experiment. One example of a trial sequence from each condition (only visual in blue, only auditory in red and audio-visual in purple. The facial images shown in this figure are AI-generated and are used for illustrative purposes only; they are not screenshots extracted from the experimental video stimuli and do not depict real individuals. **B)** Univariate responses to facial, vocal and bimodal expressions in all defined ROIs. Beta weights extracted in individually defined ROIs from independent face and voice localizers. Regions whose name is reported in blue on the x-axis are defined through the face localizer, in red through the voice localizer. The beta weights are extracted in all regions from the visual data (in blue), the auditory data (in red), and the bimodal data (in purple). Beta estimates are represented here averaged across emotion expressions. Differences from zero are evaluated through a one sample t-test, p-values are Bonferroni corrected for 12 ROIs.

In all the runs of the main experiment, participants were asked to press a button when the same emotion appeared in two events in a row, independently of the actor portraying the emotion. This ensured that their attention was on the emotional content of the expression, rather than on lower-level features of the stimulus, since the emotion repetition was always portrayed across actor identity. One third of the runs in each modality condition contained two repeated emotions (two 1-back trials), a third of the runs contained one 1-back trial while a third of the runs contained none. Each run could therefore have 20, 21 or 22 stimuli. The order of the stimuli was pseudorandomized, so that the same emotion would *never* appear twice in a row, except in the case of the planned one-back trials.

#### Acquisition parameters

Participants were scanned with a 3T GE (Signaf Premier, General Electric Company, USA, serial number: 000000210036MR03, software version: 28\LX\MR, Software release : RX28.0_R04_2020.b) MRI scanner with a 48-channel head coil at the University Hospital of St Luc, Brussels, Belgium.

In the first scanning session, a whole-brain T1-weighted anatomical scan was acquired (3D-MPRAGE; 1.0 x 1.0 mm in-plane resolution; slice thickness 1mm; no gap; inversion time = 900 ms; repetition time (TR) = 2189,12 ms; echo time (TE) = 2.96 ms; flip angle (FA) = 8°; 156 slices; field of view (FOV) = 256 x 256 mm2; matrix size= 256 X 256).

All functional runs were T2*-weighted scans acquired with Echo Planar and gradient recalled (EP/GR) imaging (2.6 x 2.6 mm in-plane resolution; slice thickness = 2.6mm; no gap; Multiband factor = 2; TR = 1750ms, TE = 30ms, 58 interleaved ascending slices; FA = 75°; FOV = 220 x 220 mm2; matrix size= 84 X 84).

### Data analyses

#### Data preprocessing and General Linear Model

The MRI data were pre-processed with bidspm (v3.0.0; https://github.com/cpp-lln-lab/bidspm; Gau et al., 2023) set up through the use of the in-house CPP_SPM repository, v0.1.0. (https://doi.org/10.5281/zenodo.3554331) and carried out through the Statistical Parametric Mapping toolbox (SPM12, RRID:SCR_007037) in Matlab, version 2016b. The preprocessing of the functional images was performed separately for each of the five tasks: the face localizer, the voice localizer, the main experiment visual, the main experiment auditory, the main experiment bimodal, and in the following order: removing of 7 dummy scans, slice timing correction, realignment, unwarping, normalization to MNI space, smoothing. Slice timing correction was performed using the 29th slice as a reference (interpolation: sinc interpolation). Functional scans from each participant were realigned and unwarped using the first image of the first run as a reference (number of degrees of freedom: 6; cost function: least square). The mean image obtained from realignment was then co-registered to the anatomical T1 image (number of degrees of freedom: 6; cost function: normalized mutual information) (Friston et al., 1995). The transformation matrix from this co-registration was then applied to all the functional images. Functional MNI normalized images were then spatially smoothed using a 3D gaussian kernel (FWHM of 6mm for the runs of the functional localizers, 2mm for the runs of the main experiment).

The anatomical T1 image was bias field corrected, segmented and normalized to MNI space (target space: IXI549Space; target space resolution: 1 mm ; interpolation: 4th degree b-spline) using a unified segmentation. The deformation field obtained from this step was then applied to all the functional images (target space resolution equal that used at acquisition; interpolation: 4th degree b-spline).

For all functional data, at the subject level, we performed a mass univariate analysis with a linear regression at each voxel of the brain, using generalized least squares with a global FAST model to account for temporal auto-correlation (Corbin et al., 2018) and a drift fit with discrete cosine transform basis (128 seconds cut-off). Image intensity scaling was done run-wide before statistical modelling such that the mean image will have mean intracerebral intensity of 100. Regressors modelled the BOLD response following the onset of the stimuli (either blocks or events) and were convolved with SPM canonical hemodynamic response function (HRF). Nuisance covariates included the 6 realignment parameters to account for residual motion artefacts.

#### Regions of interest definition

Through the face localizer contrasting face and object responses, we aimed to define the core face network (Haxby et al., 2001), including bilateral FFA, OFA and pSTS. The network could be robustly identified in most of the participants: rFFA was found in 20 out of 24 participants, lFFA and rOFA were found in 21 out of 24 participants, lOFA and rpSTS in 23 and lpSTS in all 24 participants. Additionally, The pre central gyrus (PCG) was added to the regions of interest as it robustly showed face selectivity at the group level and because this region has been previously identified as face-responsive, although not as consistently as the core face system (Rossion et al., 2012). We could identify right PCG in 22 and left PCG in 21 out of 24 participants. We could not consistently identify any other area of the extended face network, like the amygdala or the anterior temporal lobe and therefore excluded those regions from our analyses.

In order to create individual masks for each ROI, we found the activation peak within each cluster of interest (identified with the contrast faces > object; all individual peak coordinates are reported in Table S1). We then created an expanding ROI starting as a sphere and that would follow the contours of the cluster until reaching 150 voxels (this was done through the in-house CPP_ROI repository). For activation clusters that did not have a minimum of 150 voxels from the start (at a threshold of .001 uncorrected), we drew a sphere of 10mm in radius, starting at the activation peak of each ROI in each subject (but without the boundaries of the activation cluster for that region). These masks have then been used to perform multivoxel pattern analyses (MVPA), with a feature selection of 100 voxels. In three cases, even the spherical ROI mask did not allow for the inclusion of 100 voxels for MVPA; those three areas have therefore been excluded.

Analogously, with our functional voice localizer we identified the core voice network in each participant, constituted mainly by cortical territory along the mid STS/STG that has been consistently shown to preferentially respond to voice, and that from here onwards will be referred to as TVA. We could robustly identify the TVAs with our localizer: left and right TVAs were identified in all analyzed subjects. An additional peak located in the precentral gyrus (PCG) emerged through the [voice > objects] contrast and could be identified in 19 out of 24 subjects, so we included the PCG in the areas defined through the voice localizer as well. Despite the notable overlap between the visually and auditorily defined prefrontal regions, these regions show different activation profiles at the univariate level (see Figure 1D), therefore we decided to keep them separate and we will refer to the visually defined pre-central gyrus as PCGv, and to the auditorily defined one as PCGa (all individual peak coordinates are reported in Table S2). The voice localizer acquisition run of one subject had huge signal distortion in the image, so we therefore discarded that acquisition and used the group-defined clusters to build the four masks used for that subject. As for the face localizer, when the individual-activation-cluster ROIs did not reach the desired size of 150vx at a lenient threshold, we drew a sphere of 10mm in radius, starting at the individual activation peak; in two cases, even the spherical ROI mask did not allow for the inclusion of 100 voxels, so those two areas have been excluded.

All the individual masks (see Figure 1B & 1C) for both the face and the voice localizer are also shared on the OSF project.

#### Univariate analysis

In order to characterize the response profile of the defined ROIs, we extracted the mean beta estimates for each regressor in each modality condition (across all voxels and all runs of each ROI, see Figure 1D for a representation of the mean beta across regressors, by modality). For each modality of stimulation, visual (faces), auditory (voices) or audio-visual, we tested whether the mean estimate of the group was different from zero through one-sample t-tests. We additionally tested whether, separately in each ROI, the emotion regressor and the modality of stimulation predicted the beta estimate, through a generalized linear mixed model (GLMM: beta ∼ modality * emotion + (1|sub)).

Figure 2B shows the beta weights extracted from all regions for the visual (blue), auditory (red), and bimodal (purple) conditions, averaged across emotion expressions. Beta estimates for each modality, presented separately for each emotion and ROI, are provided in Supplemental Figure 1.

#### Multivariate pattern classification: within-modality

In order to tackle the question of where in the face network we can find information about facial emotions, and where in the voice network we can find information about vocal emotions we performed within-modality multivariate pattern classification. We trained an algorithm to distinguish pairs of facial expressions in each face sensitive area and pairs of vocal expressions in each voice sensitive area, and evaluated its performance (decoding accuracy, DA) in each of those ROIs. Given our 5 categories – neutral, disgust, fear, happiness and sadness – there are 10 possible pairs of expressions and the chance level for each pair is 50%. If the algorithm can distinguish pairs of expressions with an overall above chance DA, we conclude that in that ROI, information about e.g. facial emotion expressions, is represented.

This type of MVP-classification additionally allows us to answer the question of whether we can find traces of facial emotions in the voice network, and of vocal emotions in the face network.

All MVPA were implemented with a linear support vector machine, through the LIBSVM library (https://www.csie.ntu.edu.tw/~cjlin/libsvm/, (Chang & Lin, 2011), in CosmoMVPA (https://www.cosmomvpa.org, (Oosterhof et al., 2016) and Matlab, with a leave-one-run-out cross-validation procedure, and feature selection set at 100 voxels.

#### Multivariate pattern classification: across visual & auditory modalities

This analysis is implemented in ROIs that present successful within-modality decoding with both the visual and auditory stimuli. It allows us to further understand whether the emotional information in these areas are represented in a way that is, at least partly, common between the two sensory modalities, and hence, partially independent of them. This is achieved through yet another multivariate pattern classification procedure, where the training set is constituted by the data from one modality of acquisition and the test set by the data from the opposite modality. The two modalities are both used as training and testing sets, and the results are averaged. Like in the previous analysis, this decoding analysis is carried out in CosmoMVPA, through a linear support vector machine, with a leave-one-run-out cross-validation procedure and a feature selection of 100 voxels.

#### Multivariate pattern classification, between unimodal and bimodal conditions

To anticipate part of the results, the within-modality MVP-classification showed a successful decoding of emotion expressions in the bimodal condition, without however any enhancement in the performance compared to the dominant modality (aka face in the face network, voices in the voice network-see figure 2A), made us wonder whether the representation in these areas comes down to the representation of the dominant modality when this is present, without distinction between a unisensory and a bisensory context. In the light of these results, a way to explore whether a multisensory context is distinguishable from a unisensory one in these regions would be to perform MVP-classification within emotional category of different sensory contexts, specifically of the native modality of each ROI (e.g. visual facial expressions in visually-defined face network) and the bimodal condition. In order to do that, runs are taken from the bimodal condition and then either from the visual or the auditory condition; the classifier is then trained to distinguish e.g. visual disgust from bimodal disgust, visual fear from bimodal fear, and so on. If the representation of the emotional information in each ROI comes down to the dominant modality for that region, the classifier should not be able to tell the difference between e.g. auditory sadness and bimodal sadness in TVA. Importantly, although the unimodal and bimodal stimuli were presented in separate runs, this does not introduce a confound for the present analysis. The comparison between sensory contexts (e.g., visual vs. bimodal) was always performed using data from different runs for training and testing, ensuring that both conditions contributed information collected under equivalent circumstances. Any potential run-specific noise or baseline differences therefore affected the unimodal and bimodal data equally and cannot explain successful decoding. Because the classifier was trained to distinguish patterns corresponding to the same emotion across these two contexts (e.g., visual disgust vs. bimodal disgust) using independent runs for both conditions, any above-chance performance reflects genuine differences in the underlying neural representations rather than artefacts of the experimental design.

#### Brain data visualization: dissimilarity matrices, multidimensional scaling and dendrograms

To further characterize the nature of each ROI and examine in greater detail how different modalities coexist and interact within the visual and auditory networks, we focused on the neural representation of each emotion and modality. This approach allowed us to assess the similarity or differences in activity patterns across emotions and modalities.

First, for each subject, we computed a dissimilarity matrix (DSM) for each ROI, incorporating the five emotions across the three modalities. Using Spearman’s correlation, we calculated the 1 - similarity between the activity patterns for each emotion and the other emotions, both within the same modality and across different modalities. To account for potential confounds due to varying noise across runs (since each modality was presented in separate runs), we used a split-half approach. Specifically, we divided the data from each modality into two halves (i.e., odd and even runs) and averaged the activity patterns for each of the five emotions in each half. This resulted in two activity patterns for each emotion in each modality. We then computed the 1 - Spearman’s correlation between the patterns from one half and the corresponding patterns from the other half. This approach generated non-symmetrical 15x15 matrices (5 emotions x 3 modalities). To convert these matrices into symmetrical dissimilarity matrices, we averaged the values from the upper and lower triangles and set the diagonal values to zero (see Figure 4).

In addition to DSMs, we also visualize the data using 3D multidimensional scaling.

The emotions from the 3 modalities of presentation have been arranged such that their pairwise distances approximately reflect response pattern similarities. Items placed close together show similar response patterns. Items arranged far apart generated different response patterns (Mattioni et al., 2020). Finally we also visualize the brain data in the form of a dendrogram. We used MATLAB’s ‘dendrogram’ function to create a tree-like diagram that shows how the brain data is grouped into clusters. Different colors are used to highlight these clusters, based on how closely the data points are related. We used the default setting, where the colors change when the similarity between clusters drops below 70% of the highest possible value. This helps us easily see which emotions/modalities are grouped together.

#### Searchlight decoding analyses

For the decoding analyses previously described, we complemented the ROI approach with a whole-brain searchlight analysis. The primary objectives of these additional analyses were to: 1) confirm that the face and voice networks identified by the localizer were the main regions involved in the representation of facial and vocal emotions and 2) investigate whether other brain regions, outside of our functionally defined face and voice networks, were also involved in representing these emotions.

We basically repeated the three within modality decoding analyses (within visual modality, within auditory modality & within bimodal modality) and the crossmodal audio-visual decoding analysis described above, but this time the same procedure was implemented in each 100-voxels sphere using a searchlight approach (Tong & Pratte, 2012). In all the classification analyses we ran multiclass decoding analyses, testing the discriminability of patterns for the five emotions using a linear support vector machine (SVM), through the LIBSVM library (https://www.csie.ntu.edu.tw/~cjlin/libsvm/, (Chang & Lin, 2011), in CosmoMVPA (https://www.cosmomvpa.org, (Oosterhof et al., 2016) and Matlab, with a leave-one-run-out cross-validation procedure. Classification accuracy for each sphere was assigned to the central voxel of the sphere, in order to produce accuracy maps for each subject. The resulting accuracy maps were then smoothed with an 8-mm Gaussian kernel.

In a second step, we further focused on the three regions that exhibited a supramodal profile in the searchlight results—namely, the TVA, pSTS, PCG— to examine their internal spatial organization. Specifically, we aimed to investigate the presence of spatial gradients of modality sensitivity within these regions, assessing whether distinct subregions preferentially represented visual, auditory, or crossmodal emotional information. To do this, we first combined the individual ROIs from all participants to create average group-level masks for each of the six regions (left and right TVA, pSTS, and PCG). These masks were then used to constrain the searchlight results, which are displayed at an uncorrected threshold of p < 0.001 (see Figure 6).

Within these constrained regions, we examined whether subregions showed preferential decoding for visual or auditory modality, or instead exhibited crossmodal (audio–visual) integration. In the case of the TVA, we also compared the within-auditory, within-visual, and crossmodal decoding maps with the anterior, middle, and posterior TVA peaks previously identified by Pernet et al., 2015 (The exact MNI coordinates are reported in Figure 6), to assess potential correspondence with known functional subdivisions.

#### Statistical analyses

Significance of the decoding accuracies in the ROIs analyses is evaluated through a non-parametric procedure that involves permutations and bootstrapping of the data (Stelzer et al., 2013) in the following fashion: for each subject, an additional one-hundred MVP-classification procedures are performed, but where the labels for the 5 emotions are randomly assigned. Then, one value from the obtained one-hundred is sampled from each subject, ten thousand times, in order to obtain a group-level null distribution. The statistical significance of the classification results is assessed by calculating the proportion of accuracy values in the group null distribution that exceed the observed value. All p-values are then FDR corrected for multiple comparisons (number of comparisons equal to the number of ROIs tested).

For the searchlight decoding analyses we used the brain maps obtained from the searchlight analyses for each subject to run group-level statistics in SPM. Specifically, we performed a voxel-wise one-sample t-test to identify brain regions where the decoding performance was significantly above chance across participants. We report results at an uncorrected threshold of *p* < 0.005 to provide a comprehensive view of the networks involved. To ensure statistical robustness, results were also corrected for multiple comparisons using a *p* < 0.05 (FWE) threshold. These data, presented with this more stringent correction, are included in the Supplemental Information (see figure S3).

## Results

### Univariate results

The analyses of the beta estimates show that the average response in each region is significantly above zero for the stimuli in the preferred modality of each ROI, as well as for the bimodal stimuli. The FFAs and the TVAs do not show significant positive estimates for the stimulation in their non-dominant modality, vocal and facial stimulation respectively, while the OFAs show significant negative response to the voices (deactivation). The visually-defined pSTS shows significant positive response to vocal as well as facial stimuli bilaterally, with however a significantly higher response to faces, and an even higher response to bimodal stimuli. The lPCGv also shows significant positive estimates for all modalities, but without differences among them. The bilateral PCGa shows as well significant responses to all modalities, with higher estimates for the bimodal stimulation but no difference for the two unimodal ones (see Figure 2B and Supplemental Figure 1).

Finally, in only a few regions, we found a main effect of emotion (e.g., rOFA: fear > neutral; bilTVA: sadness > disgust, fear, and neutral; bilpSTS: disgust > neutral), but overall, different emotions elicited similar brain responses across all ROIs at the univariate level. Distinct responses for separate emotions were therefore mostly explored in the following multivariate analyses.

### Multivariate pattern classification, within-modality

The results of the within-modality decoding accuracies are shown in Figure 3A. We observed that emotion expressions could be robustly classified in the dominant modality of any area (facial expressions in visually defined regions and vocal expressions in auditorily defined regions, all p_FDR_ < 0.001) as well as from the bimodal conditions (all ps_FDR_ < 0.001). Additionally, vocal expressions could be selectively decoded from face-selective regions in lFFA (p_FDR_ = 0.032) and rFFA (p_FDR_ = 0.026), pSTS and PCGv, bilaterally (all ps_FDR_ < 0.001); and facial expressions could be decoded in the voice selective regions lTVA (p_FDR_ = 0.012), rTVA (p_FDR_ = 0.003) and PCGa bilaterally (ps_FDR_ < 0.001). In the FFAs and in the TVAs, the decoding accuracy is higher in their native modality than in the opposite modality, while pSTS and all the PCG (a and v) exhibit similar levels of decoding for facial and vocal expressions. In all ROIs, the decoding accuracy in the bimodal condition was not statistically different from the decoding accuracy of the dominant modality. The dominant (or native) modality of a ROI refers to the sensory modality used to define the ROI and that yields to the highest unimodal decoding accuracy, e.g. visual (facial expressions) in FFAs and audition (vocal expressions) in TVAs.

**Figure 3:**
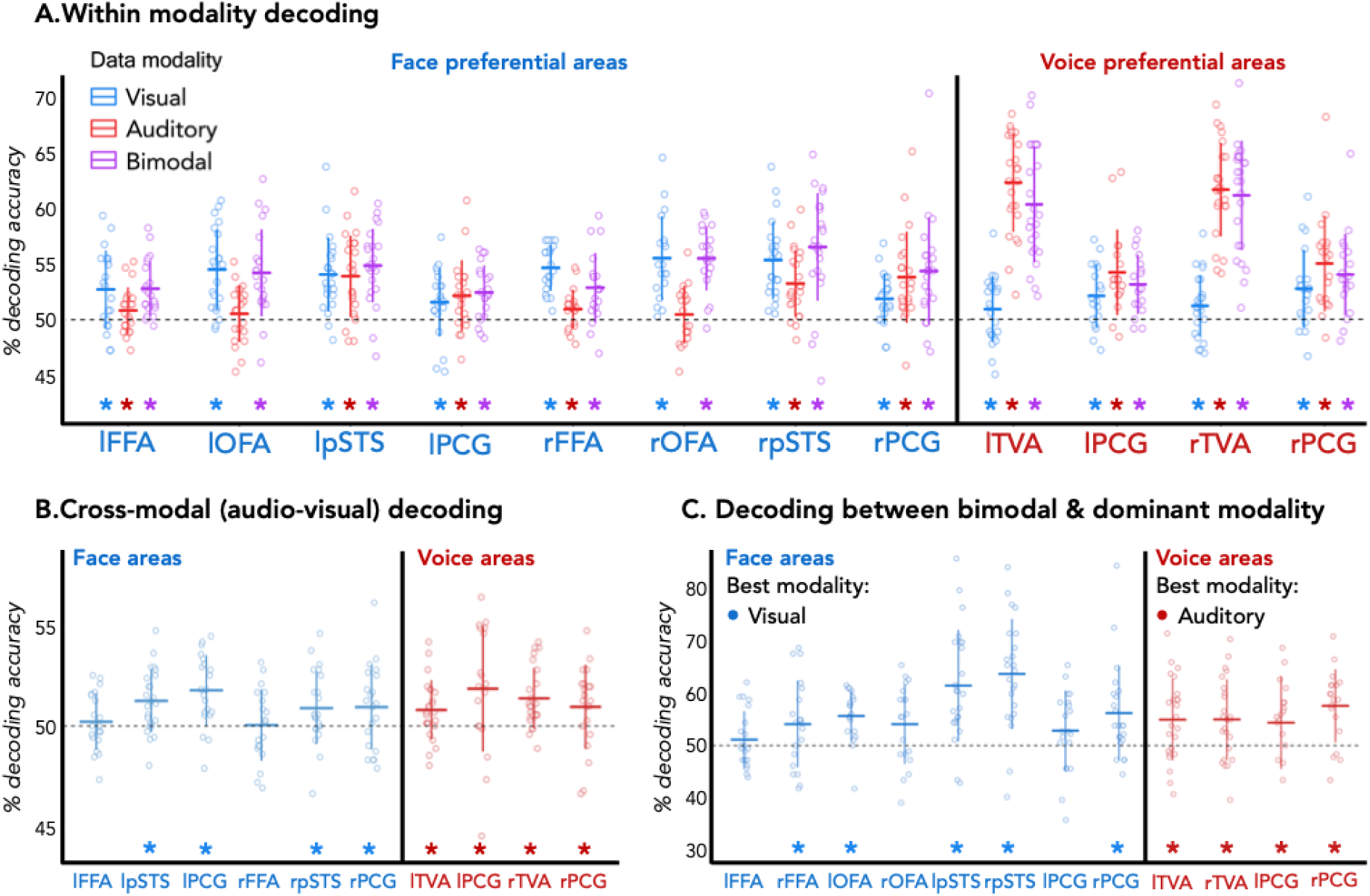
Decoding results. **A)** Within-modality classification accuracies. Regions whose name is reported in blue on the x-axis are defined through the face localizer, in red through the voice localizer. The decoding is performed in all regions with visual data (in blue), auditory data (in red), and bimodal data (in purple). Each dot represents one participant and the average of the decoding accuracies of the ten possible emotion pairs. Error bars are the SD. Significance against chance level is assessed with permutations (n = 100) + bootstrap (n = 10k), with p-values FDR corrected for 12 ROIs. **B)** Crossmodal decoding accuracies in the ten ROIS where within-modality MVPA is above chance for both vision and audition. Significance assessed with permutations (n = 100) + bootstrap (n = 10k), p-values FDR corrected for 10 ROIs. **C)** Decoding accuracies of the MVP-classification of within-emotion pairs from the data of the dominant modality and the multimodal data. In the regions identified though the face localizer (blue labels), same-emotion pairs are classified between the visual and bimodal data; in the regions identified through the voice localizer (red labels), same-emotional pairs are classified between the auditory and bimodal data. Each dot represents one subject and it is the average of the decoding accuracies of the five within-emotion pairs. Error bars are the SD. Significance against chance level is assessed with permutations (n = 100) + bootstrap (n = 10k), with p-values FDR corrected for 12 ROIs.

### Multivariate pattern classification across audio-visual modalities

We proceeded to run MVP-classification across modalities in the ROIs that showed significant within-modality decoding both for facial and vocal emotions. These areas are: bil-FFA, bil-pSTS, bil-PCGv, bil-TVA and bil-PCGa. Results are shown in Figure 3B. The pSTS, the TVA and both the PCGv and PCGa showed significant cross-modal decoding bilaterally (all ps_FDR_ ≤ 0.0015), while the crossmodal decoding accuracy in the FFAs do not differ from chance level (significance of the decoding accuracies is evaluated through the same procedure described in method section *Statistical analysis*, with FDR-correction carried out here for 10 ROIs).

### Decoding between unimodal and bimodal conditions

The within-modality MVP-classification showed a successful decoding of emotion expressions in the bimodal condition, without however any enhancement in the decoding accuracies. To follow up on these results, we performed a cross-decoding analysis between the dominant modality and the bimodal modality (e.g. visual-bimodal in FFA and auditory-bimodal in TVA) to see whether a multisensory context is distinguishable from a unisensory one in these regions. If the representation of the emotional information in each ROI comes down to the dominant modality for that region, the classifier should not be able to tell the difference between e.g. auditory sadness and bimodal sadness in TVA. The results for this analyses are shown in Figure 3C and show that overall, the classifier could distinguish unimodal vs bimodal emotions significantly in rFFA (p_FDR_ = 0.041), lOFA (p_FDR_ = 0.027), bil-pSTS (p’s_FDR_ < 0.001), rPCGv (p_FDR_ = 0.007), bil-TVA (p’s_FDR_ = 0.019), bil-PCGa (p_FDR-left_ = 0.041, p_FDR-right_ < 0.001), but only a trend is found in rOFA (p_FDR_ = 0.059), while decoding is not significantly different from chance in lFFA (p_FDR_ = 0.291), and lPCGv (p_FDR_ = 0.146). Although not completely homogeneous across regions, these findings show that there is a differential activation in the face network when voices are concurrently perceived, and the same is true in the voice network when faces are concurrently perceived; even though this never translated in a better ability of the classifier to distinguish emotion categories in the bimodal condition compared to the respective native unimodal condition in any region. These results, together with the other MVP-cassification analyses carried out in this report, speak for the multimodal nature of these networks.

### Brain data visualization: dissimilarity matrices, multidimensional scaling and dendrograms

In Figure 4, we show the group average dissimilarity matrix, multidimensional scaling, and a dendrogram for each ROI, allowing us to accurately visualize how the five categories, presented in three different modalities, cluster (or do not cluster) together. Although more qualitative in nature, data visualization provides a complementary perspective for exploring our brain data and can reveal additional nuances beyond the general results found through classification analyses.

The first key observation concerns the contribution of the non-dominant modality to emotion representation: does the bimodal condition cluster with the dominant modality, or does it form a distinct cluster? The guiding idea is that the more the bimodal representations separate from the dominant modality within a given region, the stronger the contribution of the non-dominant modality to that region’s representational space.

**Figure 4:**
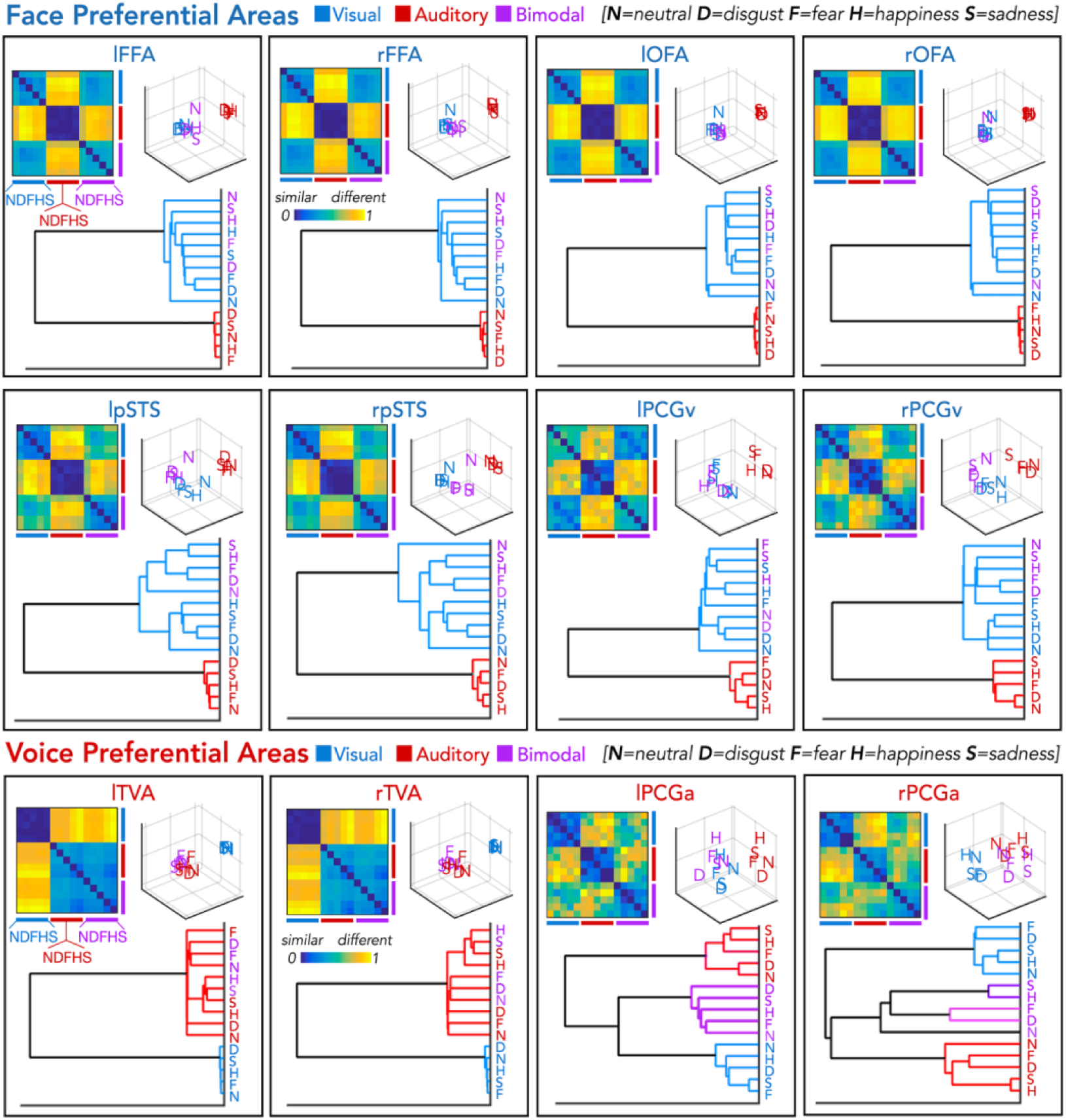
Representational similarity analyses (RSA) within face- and voice-preferential regions. Representational dissimilarity matrices (DSM), multidimensional scaling (MDS) plots, and dendrograms illustrating the neural representational structure of emotions acr oss modalities within face-preferential areas (top panels) and voice-preferential areas (bottom panels). For each region of interest (ROI), the DSMs display the pairwise dissimilarity (1 – Spearman’s r) between the neural patterns associated with the five emotion categories (N = neutral, D = disgust, F = fear, H = happiness, S = sadness) across the three modalities (visual, auditory, and bimodal). To minimize run-specific noise, a split-half approach was used (odd vs. even runs), and the resulting DSMs were averaged across the 24 subjects. In each panel, the MDS plots (3D insets) visualize the relative distances between emotion representations, where items placed closer together correspond to more similar neural patterns. Dendrograms summarize the hierarchical clustering of emotion–modality representations, with color-coded branches indicating clusters that share ≥70% similarity. Blue, red, and purple labels correspond to visual, auditory, and bimodal modalities, respectively.

The multidimensional scaling plots in Figure 4 show that in OFA the cluster from the bimodal condition overlaps with the cluster from the visual (dominant) condition, in FFA and TVA the bimodal condition start to form a distinguishable cluster from the unimodal dominant-modality and the distance between these two clusters become even more visible in the pSTS and in PCG, especially in the auditorily defined one. The cluster analysis also shows that in areas with a clear sensory preference—such as the OFA and FFA for visual processing and the TVA for auditory processing—the bimodal condition tends to cluster closely with the dominant modality. In contrast, in regions with a more multisensory profile, including the bilateral pSTS and PCG, the bimodal representations cluster less and less with the dominant modality. This pattern suggests that in these regions, the combined presentation of visual and auditory information gives rise to a unique representation distinct from those elicited by the unimodal conditions.

Interestingly, the dissimilarity matrices (DSMs; Figure 4) for the left pSTS, the right PCGv and the right PCGa—and even more prominently for the left PCGv and PCGa—show visible intermediate diagonals. This pattern indicates that representations of the same emotion (e.g., happiness) are similar across sensory modalities (visual, auditory, and bimodal). Thus, despite the three modalities maintaining distinct overall representational structures, individual emotions appear to share a common neural signature that transcends the modality of presentation.

### Searchlight decoding analyses

Results from the searchlight decoding analysis are presented in Figure 5. For decoding the five emotions within the visual modality, we observed significant decoding (p<0.005 uncorrected) in a bilateral region of the cortex, encompassing the superior and middle temporal cortex and extending to the inferior occipital cortex and to the fusiform gyrus. This network also included the bilateral postcentral gyri (PCG), inferior temporal cortex, superior and middle occipital gyri, inferior parietal cortex, supplementary motor area, medial and inferior frontal cortex, and left insula (Figure 5A).

**Figure 5:**
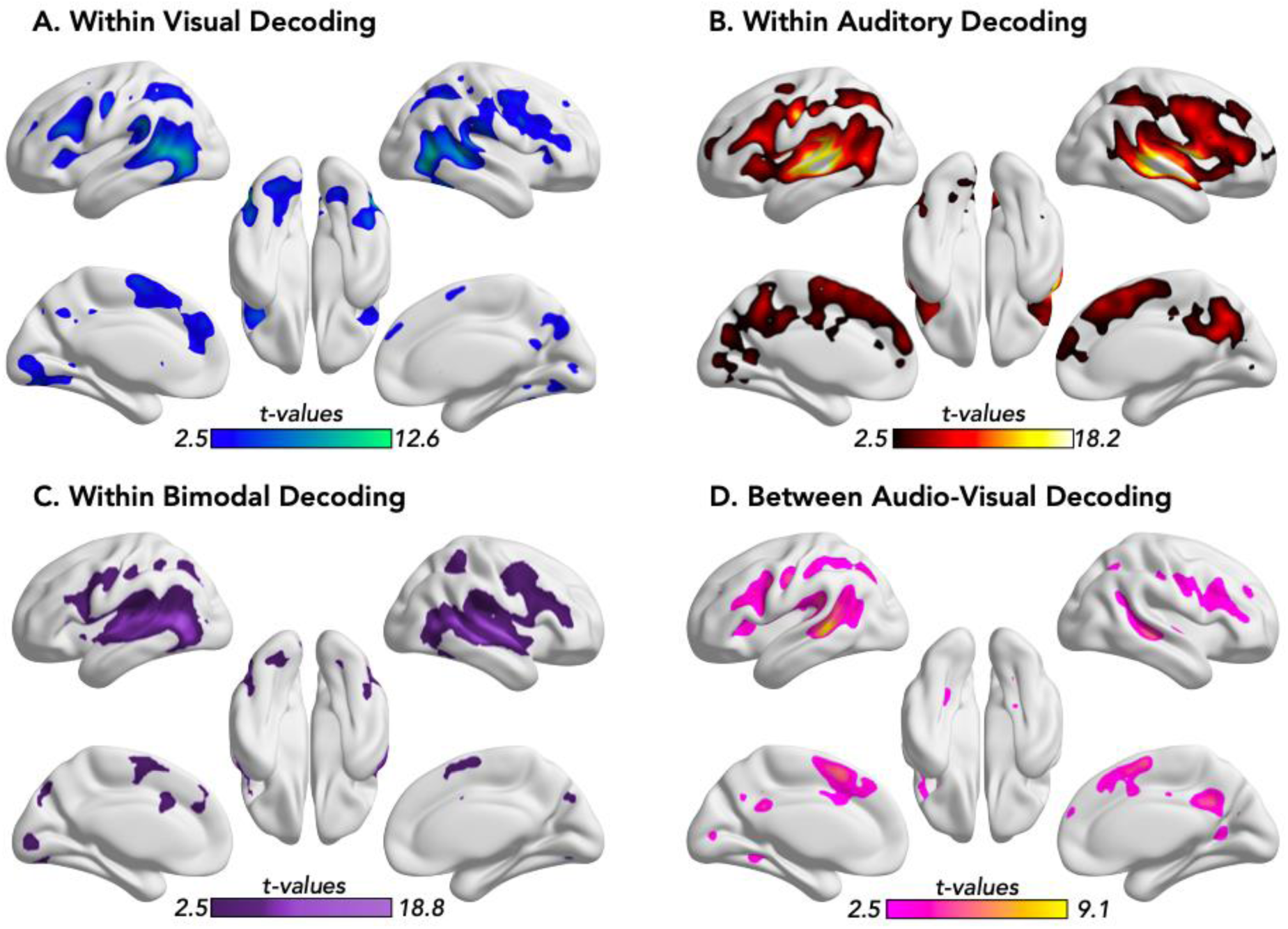
Whole-brain searchlight decoding analyses for emotion representation. Statistical maps show regions with significant decoding accuracy for emotional categories across four analyses: **(A)** within visual modality (blue/green map), **(B)** within auditory modality (red/yellow map), **(C)** within bimodal modality (dark/light purple map), and (**D)** between (crossmodal) audio–visual modalities (pink/yellow map). Each voxel represents the center of a 100-voxel searchlight sphere in which multiclass decoding was performed using a linear SVM classifier (LIBSVM; Chang & Lin, 2011) implemented in CoSMoMVPA (Oosterhof et al., 2016). Maps display group-level t-values thresholded at p < 0.005 uncorrected. Results are projected on the inflated cortical surface for visualization using BraiNet Viewer (Xia et al., 2013). Color bars indicate t-value ranges for each decoding type.

For decoding the five emotions within the auditory modality, we observed significant decoding (p<0.005, uncorrected) in the bilateral superior and middle temporal cortex, the left infero-temporal cortex, small bilateral portions of the fusiform gyrus, the bilateral postcentral gyri, the bilateral infero-parietal cortex, the bilateral medial and infero-frontal cortex and insula (Figure 5B).

For decoding the five emotions in the bimodal audio-visual condition, we observed significant decoding (p<0.005 uncorrected) in the full audiovisual network mentioned in the within-visual and within-auditory modalities, described above (Figure 4C).

Finally, for the cross-modal audio-visual decoding (Figure 4D) results, we observed bilateral middle/superior temporal cortex, the bilateral postcentral gyri, a small portion of the left fusiform gyrus, a left portion of the inferior frontal cortex, the medial frontal cortex and the left insula.

In Supplementary Figure S3, a restricted network of regions involved in within visual decoding, within auditory decoding, and between audio-visual cross-decoding is reported at a more stringent threshold (p<0.05, FEW corrected).

Beyond the whole-brain analyses, we conducted a more focused examination of the searchlight results within six regions showing a supramodal profile: the left and right TVA, pSTS, and PCG. The goal of this analysis was to assess whether these regions contain spatially organized subregions exhibiting differential sensitivity to sensory modality. In particular, we tested whether distinct subportions preferentially responded to visual or auditory information, and whether intermediate areas displayed more integrative, multimodal patterns of activation.

In the case of the TVA, we directly compared the within-auditory, within-visual, and between–audio-visual decoding results with the peaks of the anterior, middle, and posterior TVA previously identified by Pernet et al., 2015 (exact coordinates from this study are reported in figure 6). We found that the group peak for within-auditory decoding was located between the middle and anterior TVA, whereas the peak for within-visual decoding was closer to the posterior TVA. The crossmodal (audio-visual) decoding peak was situated in the region of overlap between these two areas.

**Figure 6:**
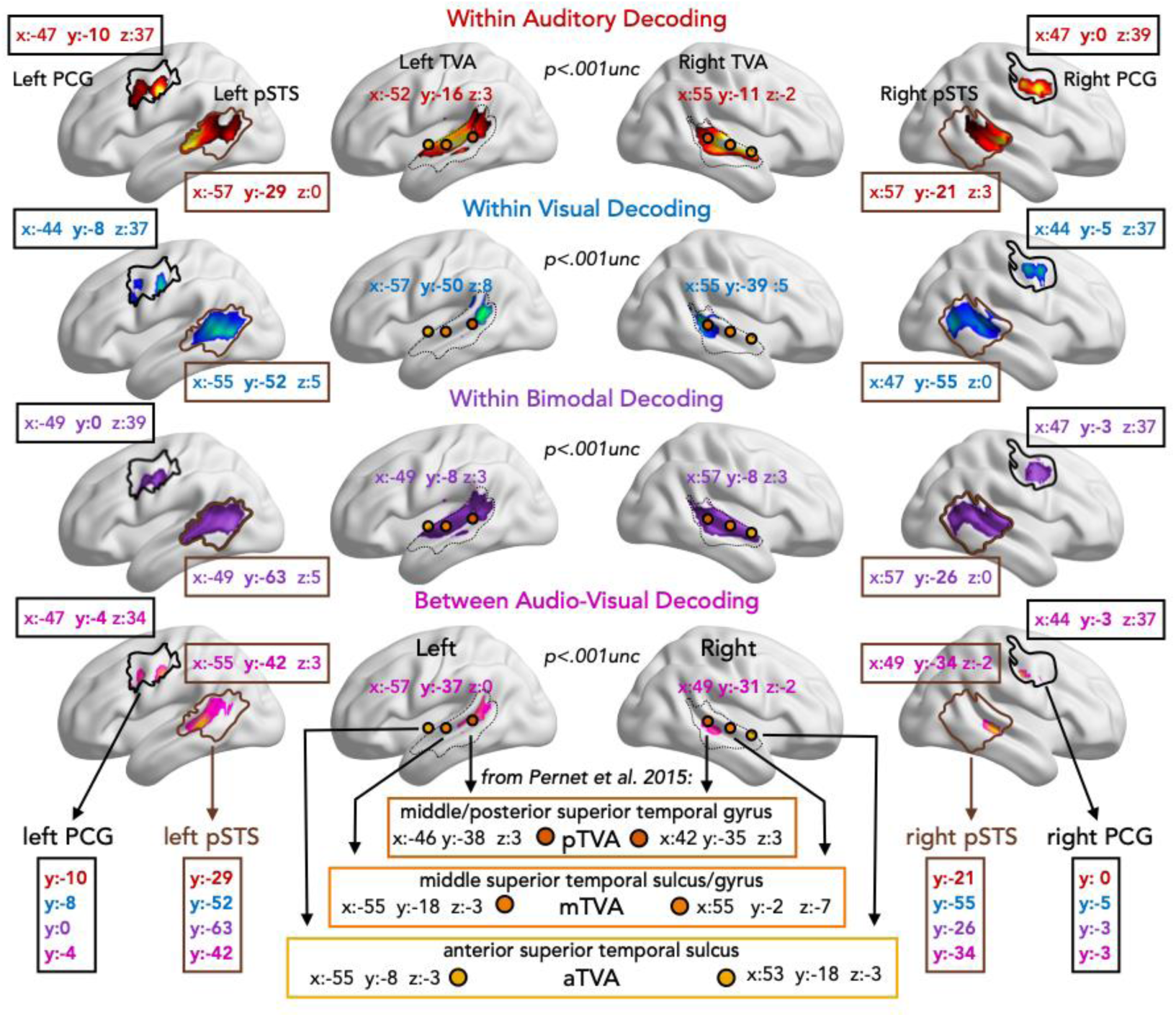
ROI-constrained searchlight decoding analyses within supramodal regions. Searchlight decoding results within the group masks of the temporal voice area (TVA), posterior superior temporal sulcus (pSTS), and precentral gyrus (PCG) for the four decoding analyses: within auditory modality (red), within visual modality (blue), within bimodal modality (purple), and between (crossmodal) audio–visual modalities (pink). Each map shows regions displaying significant decoding accuracy for emotional categories (p < 0.001 uncorrected). Reported coordinates next to each mask indicate the group decoding peaks in MNI space for each region. Orange circles with black outlines mark the anterior, middle, and posterior TVA peaks reported by Pernet et al. (2015), shown for reference to illustrate the correspondence between our crossmodal peaks and previously described TVA subregions. The orange-outlined boxes summarize the MNI coordinates of the aTVA, mTVA, and pTVA peaks from Pernet et al. (2015). Boxes at the bottom left and right of the figure summarize the group peak MNI y-coordinates for each decoding analysis, specifically along the anterior–posterior axis. Together, these results reveal a posterior-to-anterior gradient within the TVA and pSTS—posterior regions showing stronger visual decoding, anterior regions stronger auditory decoding, and crossmodal decoding emerging at their intersection—whereas the PCG exhibited overlapping peaks for all decoding types, consistent with a more integrated, supramodal representation.

A similar spatial organization was observed in both the left and right pSTS. Specifically, the within-auditory decoding peaked in the more anterior portion of the ROI, the within-visual decoding peaked in the more posterior portion, and the crossmodal decoding emerged in the intermediate region connecting the two.

In contrast, no such gradient was evident in the left or right PCG. In this region, the group peaks for the within- and between-modality decoding were spatially close, suggesting a more general modality-independent representation of emotion in this region. See Figure 6 for the MNI coordinates of the group peaks.

## Discussions

We used functional MRI to characterize the brain activity elicited by dynamic unimodal and bimodal emotional expressions conveyed through faces and voices. Using precision neuroimaging, we individually defined face- and voice-selective areas in order to spatially and functionally constrain our multivoxel decoding analyses, which aimed to determine whether these regions contain distributed information about facial and vocal emotional expressions. Our goal was to tackle four main questions. First, we investigated where facial and vocal emotional expressions are represented in the distributed voxels’ activity within each region of their respective visual and auditory networks. Second, we examined which face- and voice-selective regions represent expressions across the senses. Third, in regions showing evidence of cross-sensory emotion representations, we tested whether emotional information is represented, at least partly, independently of the sensory modality of the input. Finally, we tested whether audiovisual expressions alter the geometry of neural responses compared with unisensory stimuli.

Our results unequivocally show that emotion expressions are represented in a widespread fashion in the areas that process the respective signals through which these expressions are portrayed, i.e. the information from the emotional facial expressions is robustly represented in all eight ROIs that we independently defined with the face localizer, and the information from the vocal expressions is represented in all four ROIs that we defined with the voice localizer (Figure 3A: visual data in face-selective regions, auditory data in voice-selective regions). These results show how emotional information is represented throughout the face and the voice processing networks, and contrast with the idea that emotional facial or vocal features are only represented in some specific higher-order areas of the face and voice networks (e.g.Kriegeskorte et al., 2007). In particular, these results challenge the notion that the FFA only represents invariant information of faces like identity and gender, but not transient information like emotion expressions because the pathways involved in the processing of these two types of features would be separate (Haxby et al., 2000; O’Toole et al., 2002). Our results therefore confirm and extend previous studies showing that emotion expressions modulate the activity of the FFAs (e.g. Bernstein et al., 2018; Liang et al., 2018, p. 201; Wegrzyn et al., 2015). Secondly, we find that the face and the voice networks are not encapsulated unisensory regions immune to crossmodal emotion expressions. Specifically, we find that in all regions except the OFAs, we can classify emotional expressions even when they are delivered through their non-preferred modality (Figure 3A, vocal expressions in face-selective regions, facial expressions in voice-selective regions), showing for the first time that crossmodal responses can be reliably encoded in distributed patterns of activity in regions typically considered as purely unisensory.

Crossmodal MVP classification revealed that emotional expressions are encoded by a partially shared distributed neural code across facial and vocal modalities in multiple regions of the face and voice processing networks (Figure 3B). These areas are, bilaterally, the visually defined pSTS, the auditorily defined TVAs, and the PCG, both visually and auditorily defined.

Evidence that the face and voice networks are not impenetrable to information of different sensory modalities is corroborated by the MVP-classification of the unimodal vs the bimodal emotions (Figure 3C). Our results reveal that a multisensory setting consistently alters the distributed activity elicited by the dominant modality in face and voice networks. Even though a bimodal presentation of faces and voices did not elicit higher decoding accuracies compared to the dominant modality, the multisensory stimulation reliably affected the multivariate response to the dominant modality in nine out of the twelve investigated ROIs. These results argue against pure sensory selectivity in specific brain regions. Instead, it appears that effects are present at all levels of the networks, even in areas considered relatively lower level in the facial processing hierarchy.

To better understand how these multisensory effects are organized across the face and voice networks, we examined the representational geometry of emotional information using RSA, MDS, and clustering analyses. This complementary approach revealed that the influence of multisensory input is not uniform but follows a gradient of integration across the cortical hierarchy (Huntenburg et al., 2017). In the OFA, the bimodal condition largely overlapped with the visual (dominant) modality, whereas in the FFA and TVA, bimodal responses began to form partially distinct clusters from the unimodal dominant modality. This separation became progressively clearer in the pSTS and PCG, particularly in the auditorily defined subregions, where the bimodal representations were most distinct from the unimodal ones. These results suggest that while early, modality-specific regions primarily reflect their dominant sensory input, higher-level and more multisensory regions increasingly integrate facial and vocal emotional cues into unique, multimodal representations. These data, of course, need to be integrated with the decoding results reported above. The fact that the bimodal modality appears to cluster with the dominant modality does not prevent successful decoding of emotions between the dominant and bimodal modalities in some of these ROIs (e.g., rFFA, lOFA, and bilateral TVA). However, this suggests that the decoder is likely relying on fine-grained information within these regions to distinguish between the two conditions. In contrast, decoding may be based on more general features of the stimuli in regions with a more pronounced multisensory nature.

Our results overall seem to support the idea that the face and voice networks are less sensory-specific than once thought (Davies-Thompson et al., 2019; Peelen et al., 2010). Socially relevant information is often derived from faces and from voices, and it is therefore reasonable to think that face and voice networks interact for optimal computation and integration of those signals (Amedi et al., 2005; Froesel et al., 2022). Neuropsychological evidence supports the existence of a multimodal emotion network rather than separate, modality-specific processes, as deficits in emotion recognition are often cross-modal, affecting the recognition of both facial and vocal cues (see Kuhn et al., 2017; Young et al., 2020 for a more detailed discussion of this point). Young infants already skillfully associate socially meaningful and functionally related signals from the face and the voice (Lewkowicz & Hansen-Tift, 2012). Functional and structural connectivity studies support the existence of direct connections between the face and voice networks (Benetti et al., 2018; Blank et al., 2011; Kriegstein et al., 2005). The idea that intrinsic links exist between face and voice networks for the processing of socially meaningful stimuli resonates with studies in sensory deprived populations, where we can predict that areas for which the main sensory input is missing, would be carrying out similar computation (e.g. extraction of sensitive social information) from signals in the spared sensory modalities (Lettieri et al., 2024). The investigation of the reorganization of the face and voice networks in deprived population have indeed shown that the TVA of deaf individuals shows preferential responses for facial stimuli (Benetti et al., 2017); while in the blind population, regions typically responding to faces preferentially response to vocal information in congenitally blind people (Fairhall et al., 2017; Mattioni et al., 2020, 2022; Van Den Hurk et al., 2017). Under such an angle, we propose that face and voice sensitive regions could be intertwined functional networks involved in processing socially engaging stimuli, like emotional expressions. Even if each network prefers either facial or vocal signal, the fact that they constantly interact to solve the same computation (e.g. recognizing which emotion is expressed by a talking face) may induce crossmodal inputs to trigger activity in their non-dominant networks, often in a spatially co-registered fashion across voxels and sensory modalities as shown by our significant crossmodal decoding results. It has actually been suggested that face selectivity in the face networks (and by extension voice selectivity in the voice network) might actually arise from the preferential connectivity between those networks and the medial prefrontal cortex, a region thought to assign social values to sensory information (Powell et al., 2018). Our observation that PCG activates both for emotional face and voice and can decode emotional expressions across the senses in a spatially co-registered fashion might support this idea. Outside of the emotion domain, it has also been shown that, for example, the FFA, the OFA and the pSTS (the core face network) are implicated in responses to stimuli with social valence other than faces, like biological motion (Grossman & Blake, 2002; Sokolov et al., 2018) or when perceiving human like interactions among simple geometric shapes (Castelli et al., 2000; Schultz et al., 2003).

While MVP-classification of expressions in FFA is significant when performed with vocal emotion, this does not translate into a significant cross-modal decoding while, in contrast, the responses to facial and vocal expressions in TVA are partially shared within the same neural population. This observation might partially resonate with the enhanced decoding accuracies for facial expressions in the TVA than in FFA. One possible explanation for this imbalance relates to the fact that facial expressions are generally discriminated more reliably than vocal emotions, and that facial information tends to dominate perceptual judgments when facial and vocal expressions are incongruent (Collignon et al., 2008, 2010). It is therefore possible that the greater reliability of the facial signal constrains the extent to which facial information is represented and aligned within regions primarily processing vocal signals, which may themselves provide less reliable emotional cues. This interpretation is consistent with evidence showing that emotion recognition from voices can be modulated by unconsciously perceived facial expressions, but not by unconsciously perceived affective pictures (De Gelder et al., 2002). Further fMRI adaptation studies (Grill-Spector & Malach, 2001) may help clarify the degree of overlap between the neural populations responding to facial and vocal emotional information.

Focusing more closely on the spatial organization of the three regions showing supramodal emotion representations—TVA, pSTS and PCG—we observed a gradient of multisensory tuning (Fig. 6). Within both the TVA and pSTS, we observed a spatial organization consistent with a visual-to-auditory gradient: visual decoding peaked in more posterior portions, auditory decoding in anterior portions, and crossmodal decoding in the intermediate zone. In contrast, the PCG did not show such a gradient, as within- and between-modality decoding peaks were spatially overlapping, suggesting a more integrated representation of emotional information. Our results are consistent with the idea that the TVA comprises several bilateral patches with relatively symmetrical organization, as previously described by Pernet et al., (2015). While the precise contribution of each subregion remains debated, converging evidence suggests a functional differentiation along the anterior–posterior axis: the anterior and middle portions appear to respond more selectively to acoustic voice features (Charest et al., 2013; Giordano et al., 2014; Latinus et al., 2013), whereas more posterior areas show stronger engagement during tasks requiring audio–visual integration (Ethofer et al., 2013; Kreifelts et al., 2007, 2009; Watson et al., 2014). In line with this organization, we observed that crossmodal decoding within the TVA peaked at the transition between these anterior/middle and posterior regions, near the pSTS. This area overlaps with a segment of the posterior superior temporal sulcus previously linked to the integration of voice and visual speech information, such as lip movements (Karthik et al., 2024; Zhu & Beauchamp, 2017), reinforcing the view that posterior temporal regions form a key locus for combining auditory and visual emotional cues.

Beyond the temporal cortex, the PCG also encoded emotional information in a modality-independent manner. Unlike the TVA and pSTS, which showed a spatial gradient from auditory to visual preference, the PCG did not display such topographical organization, suggesting that it is more homogeneously multimodal in nature. Multisensory processing within frontal and prefrontal cortices has been well documented in non-human primates (Romanski, 2007; Sugihara et al., 2006), with several studies also reporting sensory convergence in the PCG. Electrophysiological recordings have shown that neurons within a restricted zone of the precentral gyrus respond to tactile, visual, and auditory stimuli (Cooke & Graziano, 2004; Graziano & Gandhi, 2000). This convergence of sensory inputs highlights the PCG as a potential site for integrating information across modalities. In line with this view, we found that the human PCG carries emotion-related information not only within but also across sensory modalities. This result converges with previous evidence implicating the PCG in the integration of emotional and motor signals (Lima Portugal et al., 2020), where activity in motor-related cortices has been interpreted as reflecting adaptive action preparation in response to emotionally salient stimuli. Our findings extend this evidence by demonstrating that the PCG, a motor-related region with established multisensory properties, encodes emotional information across modalities, maybe serving as an interface between perception, emotion, and motor preparation.

At the whole-brain level, the searchlight decoding analyses revealed a broad network of regions involved in the representation of emotion expressions both within and across modalities (Fig. 5). These results largely overlapped with our functionally localized face- and voice-selective ROIs and also extended into adjacent territories. The crossmodal searchlight results, in particular, highlighted a widespread network encompassing areas typically considered unimodal—such as the fusiform gyrus and auditory temporal cortex (including the TVA)—as well as classical multisensory hubs in the superior temporal sulcus and surrounding cortices, and extending anteriorly into temporo-frontal and prefrontal regions. Together, these findings confirm our precision neuroimaging ROI results that emotion expressions recruit a distributed, partially supramodal network that bridges sensory-specific and multimodal regions.

One potential limitation of the present study, however, concerns the possible contribution of mental imagery. Prior studies have demonstrated that imagining faces generates activity in areas of the face processing network (O’Craven & Kanwisher, 2000; Sunday et al., 2018). The possibility of auditory-induced visual imagery and of visually-induced auditory imagery cannot be fully excluded but we find it unlikely, mainly for two reasons. First, the studies that successfully elicited brain activity from imagery, achieve so by training their subjects beforehand (especially extensive training in the case of decoding analyses, see e.g. Lee et al., 2012); and by then providing specific prompts as to when and what to imagine during the experiment. Secondly and most importantly, if imagery was responsible for our successful decoding of emotions in the non-dominant modality of the examined ROIs, and subsequently of the crossmodal decoding, we would expect to find such results across all regions. This is however not the case, and of particular relevance in this regard is our decoding result in FFA: we find within-modality decoding of vocal information in the FFAs, showing that, if part of our captured activation were due to imagery, then imagery would be generating activity in FFA during auditory perception. With such a premise, we would expect successful cross-modal classification in these areas, which we do not find.

Taken together, our results suggest a distributed but still hierarchical encoding of emotion expressions across the face and voice networks. We find that both the face and the voice networks show widespread representation of emotional information, even across sensory modalities. Furthermore, a subset of the investigated areas, specifically the posterior superior temporal sulci, the temporal voice areas, and the precentral gyri, not only represent emotional information both in vision and audition, but it does so through a code that is shared between, and hence partially independent of, the sensory modalities of the input. These findings show a multisensory gradient in the representation of facial and vocal emotional expressions that converges in temporo-frontal regions where their distributed multisensory responses align to create an amodal representation of specific emotion expressions.

## CRediT Author Statement

Stefania Mattioni: Conceptualization, Methodology, Formal Analysis, Writing-Original Draft, Visualization

Federica Falagiarda: Conceptualization, Methodology, Software, Formal Analysis, Investigation, Writing-Original Draft, Visualization, Funding Acquisition

Olivier Collignon: Conceptualization, Methodology, Writing-Review and Editing, Supervision, Project Administration, Funding Acquisition

Remi Gau: Software, Data Curation Mohamed Rezk: Software

Ceren Battal: Software

Alice Van Audenhaege: Investigation

## Acknowledgements

This work was supported by the Belgian Excellence of Science program (EOS Project No. 30991544) attributed to Olivier Collignon, the Flag-ERA HBP PINT-MULTI (R.8008.19) attributed to Olivier Collignon, a mandate d’impulsion scientifique (MIS-FNRS) attributed to Olivier Collignon. Federica Falagiarda was a research fellow, Ceren Battal was a postdoctoral researcher and Olivier Collignon is a senior research associate at the National Fund for Scientific Research of Belgium (FRS-FNRS).

