## Supplemental Info for "Representation of emotional expressions across the face and voice brain networks"

### Behavioral rating of stimuli material

In addition to the two sessions that took place inside of the scanner, each participant completed a short behavioral experiment, which took place on the first day of the fMRI experiment, before the commencing of any MRI data acquisition. During this behavioral session, we asked to rate each unimodal stimulus on several scales divided into two tasks and blocked by modality (20 visual stimuli and 20 auditory stimuli rated independently, tasks' order was counterbalanced across participants). The first task consisted of an emotional rating: after the presentation of the stimulus, participants had to answer the questions of "How disgusted/fearful/happy/neutral/sad was the person in the stimulus?" on five slider scales, one per emotion. The second task consisted in answering the following question, which targeted a judgement of valence: "How positive/negative was the emotion you just perceived?", and one targeting a judgement of arousal; "How strongly did you perceive the emotion?", again on two slider scales. The behavioral rating tasks were presented in PsychoPy, the time to provide the response was self-paced/unlimited in all ratings, but the session lasted on average around 40 minutes.

These ratings have been collected with the goal of creating behavioral models to be used in further analyses that are not reported in the current manuscript.

### Face localizer and the face network

The frames composing the videos used in the face localizers were saved as individual images and then presented at the same frame rate of the original acquisition of the facial expressions (30 frames, at 29.97fps), to render them as videos on screen. The video presentation was implemented that way as image formats are typically more easily managed and give less problems than video formats when used for stimulation. Although the videos were not the exact same files, the choice of objects portrayed in the videos are taken from a subset of the objects used in Fox et al, 2009.

### Peaks of individual regions of interest for the face network

A total of 175 ROIs have been identified for the face network. The ROI masks are always centered on the individual activation peak and include a minimum of 150 voxels (voxel range of the masks: 151-271; all individual peak coordinates are reported in Table S1).

Table S1. Face network ROIs peak activation coordinates, divided by right and left hemispheres.

| | FFA | | OFA | |
|---|---|---|---|---|
| SUB | RH | LH | RH | LH |
| 001 | 46 -44 -30 | -47 -47 -26 | 23 -96 -5 | -36 -91 -15 |
| 002 | 39 -52 -15 | -44 -24 -21 | 44 -73 -8 | -36 -91 -15 |
| 003 | no peak | -42 -34 -15 | 21 -94 5 | -29 -94 -8 |
| 004 | 39 -55 -18 | -39 -47 -23 | 34 -96 -2 | -29 -89 -18 |
| 005 | 47 -47 -21 | -47 -60 -15 | 42 -73 -8 | -47 -73 -21 |
| 006 | 30 -52 -15 | -36 -63 -15 | 26 -91 -10 | -21 -96 -15 |
| 007 | 39 -37 -21 | -44 -47 -21 | 23 -89 -18 | -23 -91 -15 |
| 008 | 47 -53 -28 | -44 -50 -26 | 28 -94 -6 | -44 -86 -10 |
| 009 | 42 -57 -23 | -42 -39 -21 | 36 -68 -5 | -36 -81 -8 |
| 010 | 47 -52 -21 | -44 -44 -23 | 49 -73 -10 | -42 -68 -10 |
| 011 | no peak | no peak | 44 -78 -10 | -23 -96 0 |
| 012 | 39 -52 -21 | no peak | 23 -94 -5 | -31 -91 -13 |
| 013 | 36 -47 -23 | -42 -47 -23 | 39 -81 -10 | -49 -76 -15 |
| 014 | 39 -52 -26 | -44 -50 -31 | 49 -73 -13 | -34 -89 -2 |
| 015 | no peak | -42 -63 -10 | 34 -91 -21 | -36 -86 -21 |
| 016 | 39 -52 -23 | -42 -42 -26 | 49 -76 -10 | -42 -86 -10 |
| 017 | 39 -39 -21 | -36 -54 -15 | 29 -96 -5 | -44 -83 -13 |
| 018 | 39 -60 -18 | -36 -65 -15 | 39 -81 -10 | -23 -86 -8 |
| 019 | 42 -47 -18 | no peak | 21 -99 -5 | -42 -86 -10 |
| 020 | 42 -47 -23 | -36 -47 -23 | no peak | no peak |
| 021 | 44 -42 -15 | -36 -60 -10 | no peak | 49 -70 -10 |
| 022 | 47 -42 -23 | -39 -42 -23 | 39 -86 -15 | -42 -83 -15 |
| 023 | 42 -57 -13 | -47 -60 -23 | 42 -73 -15 | -26 -99 -13 |
| 024 | no peak | -47 -65 -18 | no peak | -29 -94 -13 |

| | pSTS | | PCG | |
|---|---|---|---|---|
| SUB | RH | LH | RH | LH |
| 001 | 62 -42 -5 | -62 -63 -2 | 42 2 60 | -44 -5 50 |
| 002 | 52 -60 11 | -52 -52 13 | -47 -5 47 | 47 -5 60 |
| 003 | 65 -44 8 | -60 -47 13 | 47 5 50 | -42 -3 60 |
| 004 | 60 -55 0 | -62 -60 13 | 55 5 44 | no peak |
| 005 | 49 -55 13 | -42 -63 16 | 47 2 50 | 47 2 50 |
| 006 | 55 -60 8 | -55 -50 13 | 49 8 50 | -47 2 52 |
| 007 | 44 -42 8 | -47 -52 11 | 49 0 47 | -39 -3 65 |
| 008 | 44 -52 8 | -49 -55 8 | 55 0 44 | -39 -3 47 |
| 009 | 62 -34 5 | -55 -47 8 | 49 5 47 | -47 0 47 |
| 010 | 57 -29 5 | -47 -65 13 | 49 5 47 | -44 5 44 |
| 011 | 55 -47 5 | -60 -60 13 | 49 5 47 | -42 -5 50 |
| 012 | 62 -52 5 | -55 -44 13 | no peak | no peak |
| 013 | 60 -50 -5 | -60 -63 13 | 47 5 55 | -47 0 52 |
| 014 | 55 -52 5 | -62 -47 5 | 55 8 44 | -49 0 50 |
| 015 | 49 -34 -2 | -49 -39 0 | 52 8 42 | -42 2 47 |
| 016 | 60 -63 5 | -55 -70 13 | 49 0 52 | -49 0 50 |
| 017 | 47 -52 5 | -60 -52 3 | 55 5 44 | -55 15 42 |
| 018 | 52 -63 11 | -60 -47 8 | 42 2 52 | -39 -8 55 |
| 019 | 47 -34 3 | -55 -39 5 | 49 10 47 | -44 0 57 |
| 020 | 47 -55 5 | -44 -57 8 | 52 2 47 | -47 -5 42 |
| 021 | 57 -44 8 | -44 -55 11 | 55 0 52 | -39 -8 55 |
| 022 | 52 -44 5 | -49 -44 3 | 44 -3 44 | -47 2 47 |
| 023 | 52 -42 8 | -57 -50 13 | 49 2 52 | -57 -5 47 |
| 024 | no peak | -60 -47 5 | no peak | -44 5 55 |

Masks typically have a number of voxels that is slightly higher than the number selected for MVP-classification, because these areas, located at the edge of the brain and defined through a different acquisition run (the face localizer), often do not overlap 100% with activation data, i.e. not all voxels in the mask are filled with active brain voxels from the main experiment acquisitions used for the multivariate analysis. Additionally, we allowed the spherical ROIs to have a higher number of voxels (a 10mm radius sphere with 2.6mm isotropic voxels includes ~238vx) than the average mask created with the former method because the proportion of "void voxels" is higher for spheres than for a ROI defined from the individual activation cluster of the localizer map of each subject.

#### Voice localizer and the voice network

The audio files used in the "auditory objects" blocks of the functional voice localizer run are copyright free audio clips downloaded from the internet and that have been cut to a one-second duration, in order to match the duration of the voice tracks. The list of auditory objects is the following: an applause, a bike bell, a church bell, cutting scissors, a cracking egg, an engine turning on, a grinder, a hairdryer, typing on a keyboard, bowl mixing, opening can, phone ringing, streaming water, a manual saw, an electric sharpener, a thunder, toothbrushing, traffic sound, water pouring and wind sound.

#### Peaks of individual regions of interest for the voice network

There was a technical problem during the acquisition of the voice localizer of subject number 003, therefore individually-defined peaks are missing for this participant and masks created with group-level coordinates have been used (binary group masks were built through the same procedure as individual ones). For the voice localizer, a total of 84 ROI masks were used across 24 subjects. The masks are always centered on the individual highest activation peak (except in the case of sub-003) and include a minimum of 150 voxels (voxel range: 151-229; all individual peak coordinates are reported in Table S2).

Table S2. Voice network ROIs peak activation coordinates

| | TVA | | PCG | |
|---|---|---|---|---|
| SUB | RH | LH | RH | LH |
| 001 | 65 -5 0 | -68 -29 11 | 52 -11 42 | -52 -11 44 |
| 002 | 68 -31 0 | -65 -5 3 | 47 -8 57 | -49 -8 52 |
| 003 | - | - | - | - |
| 004 | 57 -24 -5 | -68 -34 5 | no peak | no peak |
| 005 | 62 -18 -5 | -57 -11 3 | 52 -3 47 | -49 -3 50 |
| 006 | 62 -31 3 | -65 -34 11 | no peak | no peak |
| 007 | 52 -29 3 | -68 -21 8 | 55 0 47 | -44 -5 55 |
| 008 | 47 -39 11 | -57 -44 16 | 57 0 44 | -55 -3 50 |
| 009 | 70 -18 13 | -68 -29 11 | 55 -3 55 | -49 -3 50 |
| 010 | 65 0 -2 | -60 -18 11 | 44 2 47 | -52 -5 47 |
| 011 | 55 -31 8 | -60 -26 8 | 49 0 47 | -44 8 34 |
| 012 | 70 -16 3 | -65 -26 5 | 49 2 52 | -49 -3 52 |
| 013 | 49 -31 5 | -65 -8 -2 | 47 -5 44 | -47 5 55 |
| 014 | 60 -11 -5 | -60 -8 -5 | 55 5 42 | -47 -5 52 |
| 015 | 65 -13 3 | -62 -16 0 | 57 2 42 | -42 2 47 |
| 016 | 60 -5 -8 | -62 -3 -5 | 55 5 44 | -52 -5 47 |
| 017 | 57 -24 -5 | -65 -11 0 | 55 0 52 | -52 -3 55 |
| 018 | 62 -3 -8 | -52 -13 5 | 52 -5 44 | -52 -8 52 |
| 019 | 62 -24 -5 | -68 -18 -2 | 49 5 47 | -49 5 44 |
| 020 | 60 -13 5 | -55 -13 5 | 55 0 42 | -52 -11 47 |
| 021 | 62 -16 -5 | -62 -18 -2 | no peak | no peak |
| 022 | 60 0 -5 | -57 -13 0 | 52 0 50 | -55 -5 47 |
| 023 | 60 -29 -5 | -65 -31 5 | 49 0 50 | -44 0 55 |
| 024 | 62 -16 0 | -65 -16 -2 | no peak | no peak |

Univariate results from event-related experiment

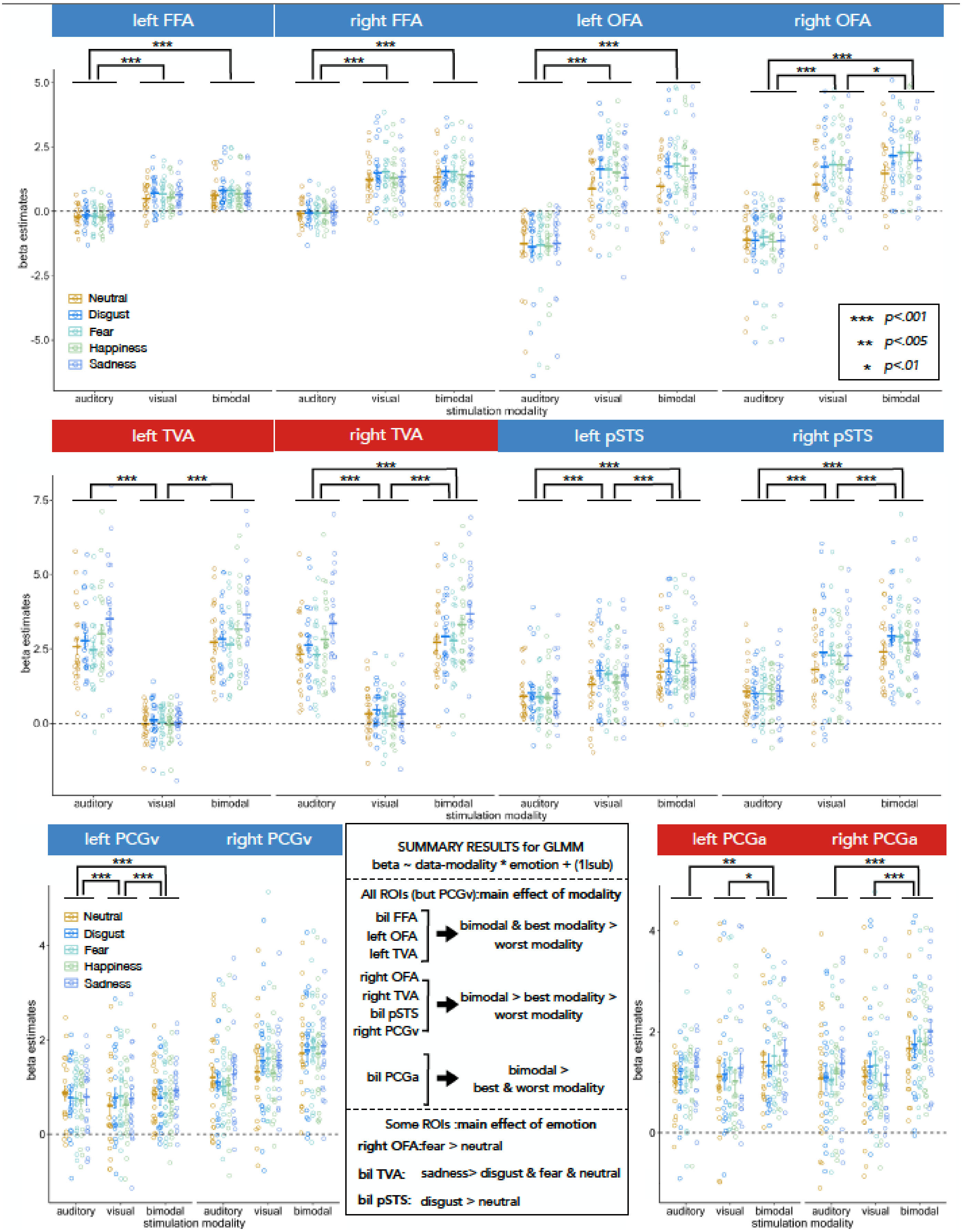


**Fig S1. Univariate results for the event-related experiment.** Average beta estimates are shown for each emotion (neutral, disgust, fear, happiness, sadness) and for each modality of stimulation (visual, auditory, bimodal) across all regions of interest (ROIs; blue labels indicate visually defined ROIs, red labels indicate auditorily defined ROIs). Bars represent mean beta values ± SEM. Stars indicate significant differences according to the mixed-effects model (β ~ data-modality × emotion + (1 | sub)): *** $p < 0.001$; ** $p < 0.005$; * $p < 0.01$. Summary effects (bottom panel) show that all ROIs except PCGv exhibited a main effect of modality. Specifically, bilateral FFA, left OFA, and left TVA showed higher activation for the bimodal and dominant modalities compared to the non-dominant one; right OFA, right TVA, bilateral pSTS, and right PCGv displayed a graded pattern with bimodal > dominant > non-dominant responses; and bilateral PCGa showed stronger responses for the bimodal than for both unimodal conditions. In addition, some regions exhibited a main effect of emotion: right OFA (fear > neutral), bilateral TVA (sadness > disgust, fear, and neutral), and bilateral pSTS (disgust > neutral).

### MVPA and permutation tests

The main report of the current study reports the significance levels of the MVPA analysis obtained through the bootstrap (N=10000) of a permutation test. The percentage of correct responses as average of the binary decoding, as well as the uncorrected and FDR-corrected p-values of the permutation tests are reported in the following tables. Table S3 reports the within-modality MVPA results, while table S4 reports the crossmodal-decoding results.

Table S3. P-values of permutation tests results – within-modality decoding. Asterisks next to the percentage decoding indicates a p<0.05 after FDR-correction (Corr. column).

| | VISUAL | | | AUDITORY | | | BIMODAL | | |
|---|---|---|---|---|---|---|---|---|---|
| ROI | %(dec) | Unc. | Corr. | %(dec) | Unc. | Corr. | %(dec) | Unc. | Corr. |
| rFFA | 54.7* | 0.0001 | 0.00013 | 51* | 0.0193 | 0.02573 | 52.9* | 0.0001 | 0.0001 |
| rOFA | 55.6* | 0.0001 | 0.00013 | 50.5 | 0.1591 | 0.1591 | 54.3* | 0.0001 | 0.0001 |
| rpSTS | 55.4* | 0.0001 | 0.00013 | 53.3* | 0.0001 | 0.00015 | 53.3* | 0.0001 | 0.0001 |
| rPCG | 51.9* | 0.0001 | 0.00013 | 53.9* | 0.0001 | 0.00015 | 54.4* | 0.0001 | 0.0001 |
| lFFA | 52.7* | 0.0001 | 0.00013 | 50.8* | 0.027 | 0.0324 | 52.8* | 0.0001 | 0.0001 |
| lOFA | 54.6* | 0.0001 | 0.00013 | 50.5 | 0.097 | 0.1058 | 54.6* | 0.0001 | 0.0001 |
| lpSTS | 54.1* | 0.0001 | 0.00013 | 53.9* | 0.0001 | 0.00015 | 54.9* | 0.0001 | 0.0001 |
| lPCG | 51.6* | 0.0004 | 0.00048 | 52.2* | 0.0001 | 0.00015 | 52.2* | 0.0001 | 0.0001 |

| | | | | | | | | | |
|---|---|---|---|---|---|---|---|---|---|
| rTVA | 50.9* | 0.0027 | 0.00295 | 61.7* | 0.0001 | 0.00015 | 61.2* | 0.0001 | 0.0001 |
| rPCG | 52.8* | 0.0001 | 0.00013 | 55.1* | 0.0001 | 0.00015 | 54.1* | 0.0001 | 0.0001 |
| lTVA | 51.2* | 0.0117 | 0.0117 | 62.4* | 0.0001 | 0.00015 | 60.4* | 0.0001 | 0.0001 |
| lPCG | 52.1* | 0.0001 | 0.00013 | 54.3* | 0.0001 | 0.00015 | 53.2* | 0.0001 | 0.0001 |

Table S4. Permutation test results – crossmodal audio-visual decoding. Asterisks next to the percentage decoding indicates a p<0.05 after FDR-correction (Corr. column).

| ROI | %(dec) | Unc. | Corr. |
|---|---|---|---|
| rFFA | 50.1 | 0.4182 | 0.4182 |
| rpSTS | 50.9* | 0.0002 | 0.00033 |
| rPCG | 50.9* | 0.0006 | 0.00086 |
| lFFA | 50.2 | 0.1784 | 0.1983 |
| lpSTS | 51.4* | 0.0001 | 0.00013 |
| lPCG | 51.8* | 0.0004 | 0.00048 |
| rTVA | 51.4* | 0.0001 | 0.00025 |
| rPCG | 51* | 0.0012 | 0.0015 |
| lTVA | 50.8* | 0.0002 | 0.00033 |
| lPCG | 51.8* | 0.0001 | 0.00025 |

### Control analysis : the role of identity

In order to rule out the possibility that a few of the utilized exemplars among the 20 stimuli could drive the emotion decoding results, an additional analysis has been carried out. We wanted to address the potential concern that the above chance classification results (main figure 3) might not be due to the recognition of the emotions in question, but rather to a few particularly deviant tokens that would be allegedly easy to single out and pair with the expected emotional label.

If a few of the utilized stimuli were easier for participants to single out while perceiving them, e.g. particularly goofy in their emotion portrayal, and if that had ended up affecting the classification of emotion regressors - despite such regressors being modeled with the contribution of all the exemplars for each emotion - a classifier should be able to more easily distinguish these stimuli compared to the others. The approach we have taken is therefore still that of multivariate pattern classification: we have performed within-modality MVPA, with the visual data in visually defined-areas, and auditory data in auditorily-defined areas, and trained the classifier to distinguish each pair of stimuli, for the total of the 190 possible stimuli pairs. The "distinguishability" of each stimulus is then expressed as the average decoding accuracy of the 19 binary pairs including the stimulus in question. To test whether the distinguishability of any particular stimuli would stand out, the average decoding accuracies are entered in a generalized linear mixed model, per ROI (GLMM with the following formula: *decoding accuracy ~ stimulus +(1|subj)*). The global effect of the *stimulus* predictor is evaluated through an analysis of variance (Kenward-Rogers degrees of freedom approximation method).

The results show a non-significant effect of the stimulus predictor in nine ROIs ($F_{lFFA} = 0.757$, $p_{lFFA} = 0.756$; $F_{rFFA} = 1.34$, $p_{rFFA} = 0.155$; $F_{lOFA} = 0.82$, $p_{lOFA} = 0.683$; $F_{rOFA} = 1.46$, $p_{rOFA} = 0.101$; $F_{lpSTS} = 1.193$, $p_{lpSTS} = 0.259$; $F_{lPCGv} = 0.631$, $p_{lPCGv} = 0.882$; $F_{rPCGv} = 1$, $p_{rPCGv} = 0.455$; $F_{lPCGa} = 1.275$, $p_{lPCGa} = 0.197$; $F_{rPCGa} = 1.194$, $p_{rPCGa} = 0.26$), demonstrating a fully comparable performance of the classifier for all twenty stimuli. The ROIs where the main effect of stimulus is significant are the lTVA ($F = 3.603$; $p < 0.001$ ), the rTVA ($F = 3.219$ ; $p < 0.001$ ) and the rpSTS ($F = 2.669$; $p < 0.001$ ), here we find that a few stimuli are significantly different than a few others (p's of post-hoc pairwise comparisons corrected for a family of 20 estimates). Specifically, in left TVA we find the decoding accuracy for stimulus number 10 to be different from stimuli 1, 3, 5, 6, 9, 11, 12, 16 and 19 (all p's $< 0.048$), additionally stimulus 3 and 17 are significantly different from each other ($p = 0.046$). In the right TVA we find stimulus 18 to be significantly different from stimuli 2, 5, 6, 9, 11, 15, 16 (all p's $< 0.049$), and stimulus 6 additionally from stimuli 8, 10 and 20 (all p's $< 0.03$). Lastly, in the right pSTS, stimulus 1 is different from stimuli 8, 13, 15, 16, 17 and 19 (all p's $< 0.039$).

Overall, these results show how there is no pattern of deviant distinguishability relating to some of the stimuli across ROIs. In fact, partly to our surprise, no stimulus stands out in the vast majority of the investigated regions. For the three regions that show some difference among stimuli, no consistency is shown across results, ruling out the possibility that any of the utilized stimuli is perceived as deviant with any consistency across subjects. Had we had found any particular stimulus, or subset of stimuli, to be standing out consistently across regions, it would have been interesting to see whether those stimuli belonged to the same identity, i.e. same actor portraying the emotions. Since this has not been the case, however, we feel confident in ruling out the potential influence of a specific identity or a specific subset of stimuli in the emotion classification analyses. The decoding accuracies by native-modality stimulus in each ROI are represented in figure S2.

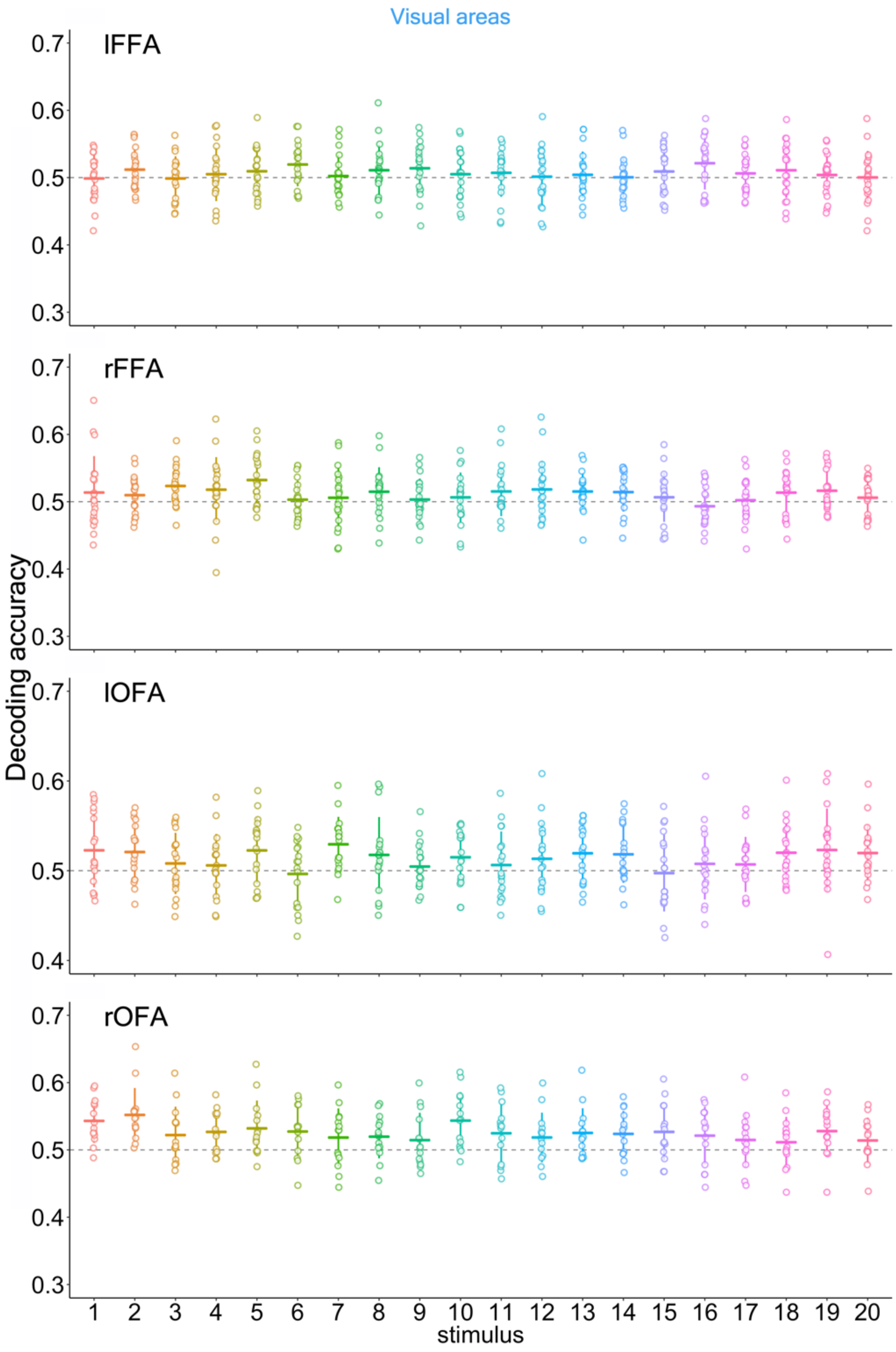


Fig. S2. Continues in the following page.

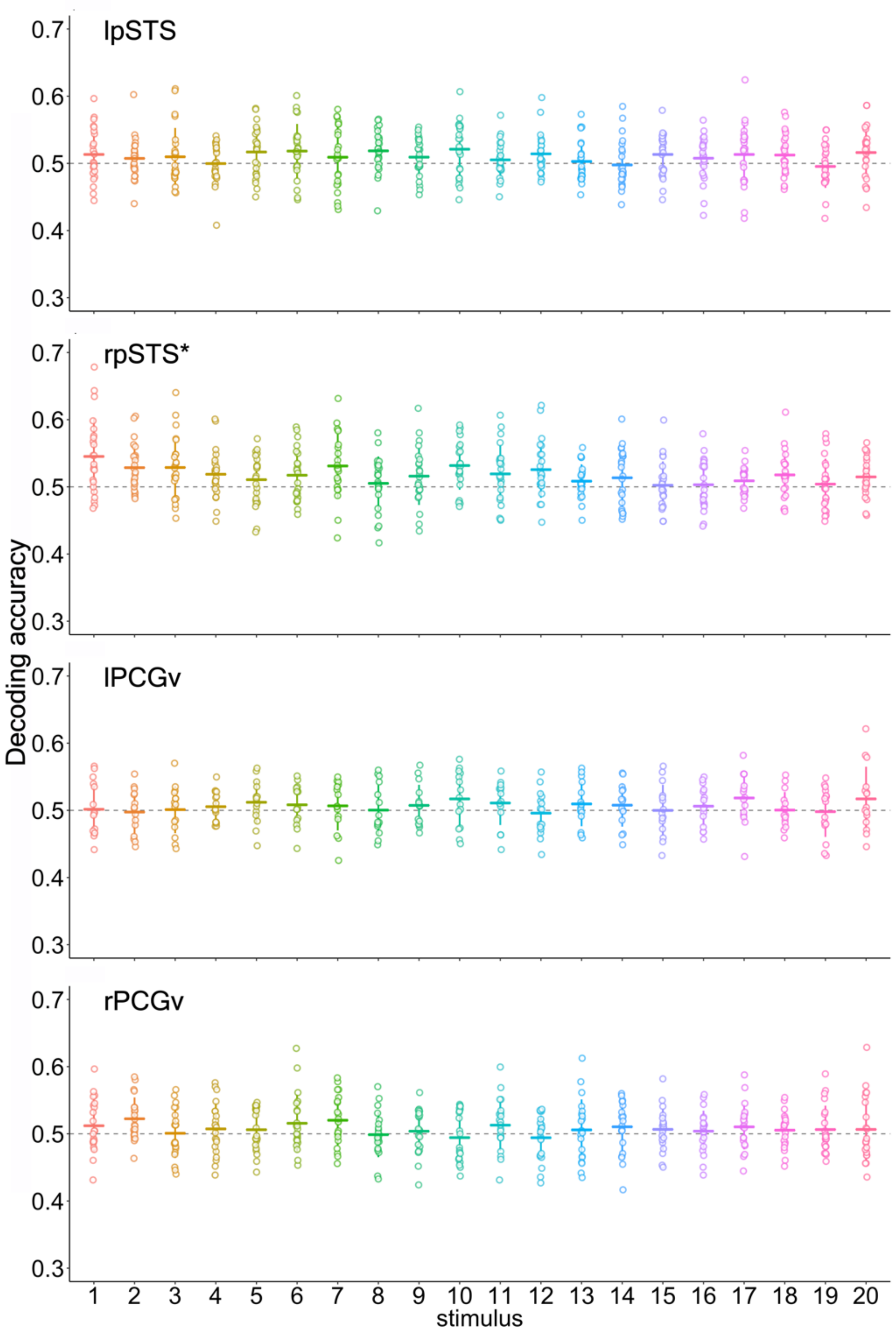


Fig. S2. Continues in the following page.

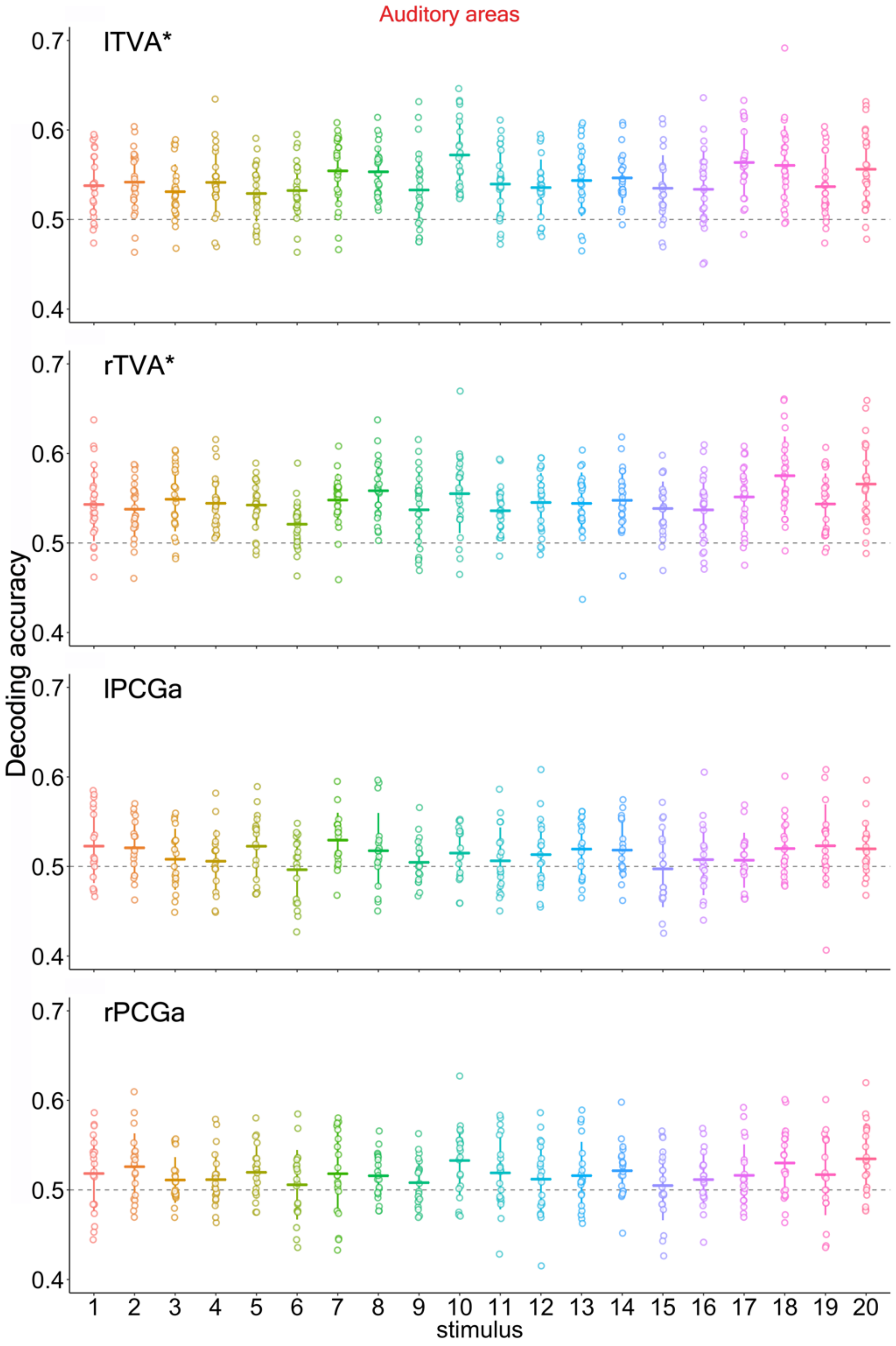


Fig S2.

Each graph shows the decoding accuracy (DA) for each stimulus (as the average of the 19 pairs in which that stimulus is present, within modality for the native modality of each area). A DA equal to chance level indicates that the specific stimulus was not distinguishable, on average, from the other 19 stimuli within the same modality. Error bars are the SD. A asterisk next to the ROI name indicates a significant effect of the factor *stimulus*. (see written paragraph for details on which decoding accuracy is significantly different from others).

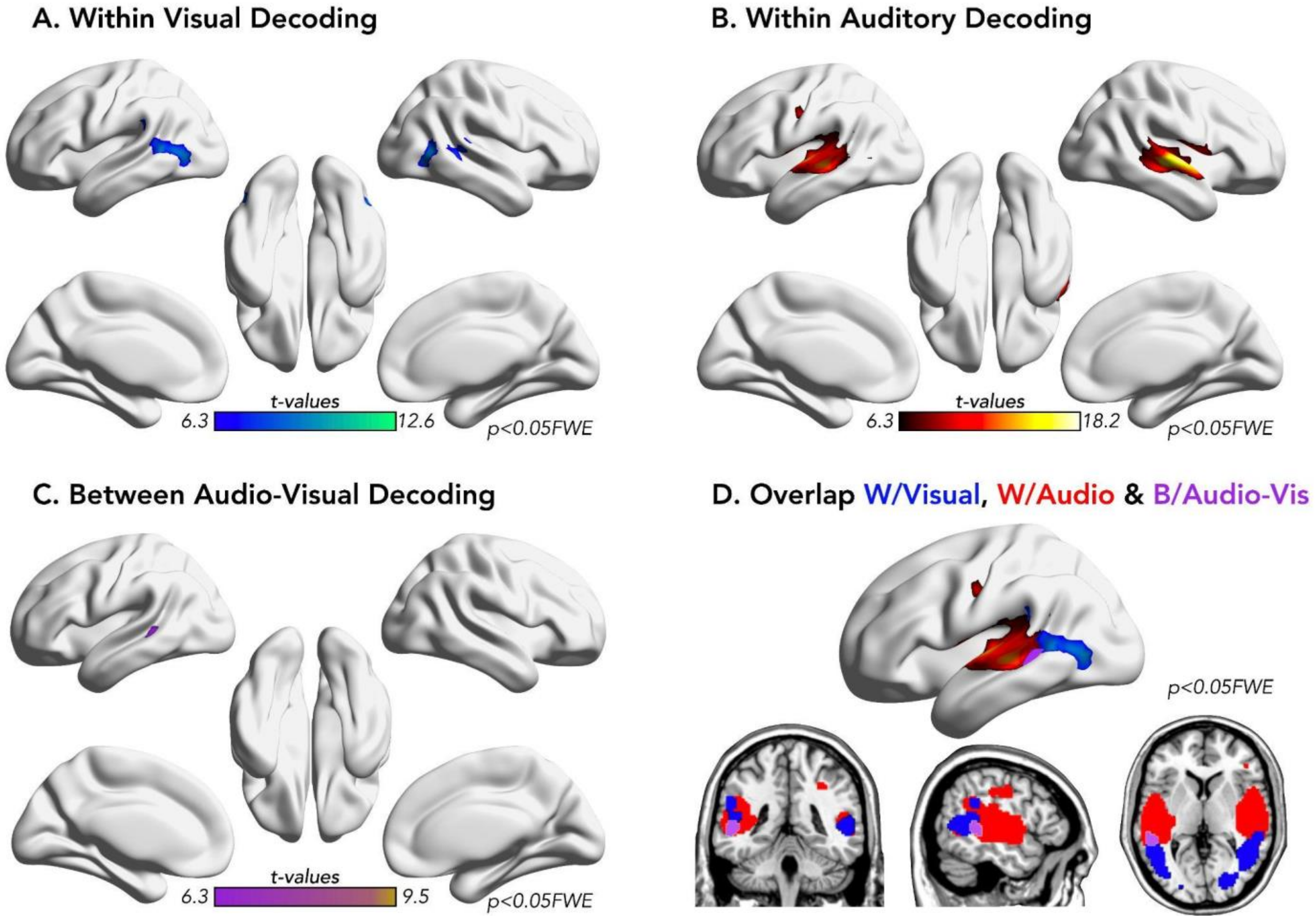


Fig S3.

Whole-brain searchlight decoding analyses for emotion representation at a stringent threshold of p<0.05FWE corrected. Statistical maps show regions with significant ($p<0.05$ FWE) decoding accuracy for emotional categories across four analyses: **(A)** within visual modality (blue/green map), **(B)** within auditory modality (red/yellow map), **(C)** within bimodal modality (dark/light purple map), and (**D**) the overlap between the three. Each voxel represents the center of a 100-voxel searchlight sphere in which multiclass decoding was performed using a linear SVM classifier (LIBSVM; Chang & Lin, 2011) implemented in CoSMoMVPA (Oosterhof et al., 2016). Maps display group-level t-values thresholded at $p < 0.05$ FWE corrected. Results in paanles A,B, and C are projected on the inflated cortical surface for visualization using BraiNet Viewer (Xia et al., 2013). Color bars indicate t-value ranges for each decoding type.
